# Efferent projections of topographically distinct relaxin family peptide receptor-3 (RXFP3) lateral hypothalamus/zona incerta cells

**DOI:** 10.64898/2026.01.15.699820

**Authors:** Brandon K Richards, Alexander IJ Kilby, Jennifer L Cornish, Jee Hyun Kim, Andrew J Lawrence, Christina J Perry

**Affiliations:** School of Psychological Sciences, Macquarie University, North Ryde, NSW 2113, Australia; IMPACT – The Institute for Mental and Physical Health and Clinical Translation, School of Medicine, Deakin University, Geelong, VIC 3220, Australia; The Florey Institute of Neuroscience and Mental Health, Parkville, Melbourne, VIC 3052 Australia; Florey Department of Neuroscience and Mental Health, The University of Melbourne, Parkville, VIC 3052, Australia

**Keywords:** RXFP3, zona incerta, lateral hypothalamus, anterograde tracing, neuroanatomy

## Abstract

Relaxin family peptide receptor-3 (RXFP3) is a ligand-activated G-protein coupled receptor and the cognate receptor for the conserved neuropeptide relaxin-3. We recently demonstrated that chemogenetically activating an RXFP3-expressing population in the lateral hypothalamus and zona incerta (LH/ZI^RXFP3^) induced escape-like jumping behaviour following fear conditioning, but only in a subset of mice. Given the diverse hodology of the LH and ZI, we hypothesised that LH/ZI^RXFP3^ cells may consist of discrete subpopulations with unique connectivity patterns that govern different aspects of defensive behaviour. To explore this possibility, we unilaterally injected small volumes of a Cre-dependent anterograde viral tracer into four distinct sites of the LH/ZI in RXFP3-Cre mice and analysed their brain-wide efferent connectivity patterns. Each injection site group produced unique projection patterns, particularly to nuclei involved in threat and defensive behaviour. Of note were strong projections from the rostral ZI and anterior LH to the lateral habenula, and projections from the intermediate and caudal ZI to the ventrolateral periaqueductal gray. By combining retrograde tracing and RNAscope fluorescent *in situ* hybridisation, we identified that most LH/ZI^RXFP3^ projections to the lateral habenula arose from a subset of vGlut2-expressing lateral hypothalamus neurons, while most projections to the ventrolateral periaqueductal gray arose from a subset of GAD1-expressing zona incerta neurons. Taken together, our results strongly suggest that LH/ZI^RXFP3^ cells exhibit distinct efferent projection patterns throughout the brain depending on their topographical location within these nuclei, likely reflecting the functional diversity of these neurons.

## 1. Introduction

The zona incerta (ZI) is a predominantly GABAergic subthalamic nucleus, consisting of several neurochemically distinct but heterogeneously organised subpopulations with extensive input-output connectivity patterns (Arena et al., 2024; Mitrofanis, 2005). Because of this, it is unsurprising that a unified definition of its function has remained elusive since it was coined as the ‘zone of uncertainty’ by Forel (1877). Indeed, the ZI has been implicated in behaviours spanning numerous functional domains, including sleep (Blanco-Centurion et al., 2023; Vidal-Ortiz et al., 2024; Zhu et al., 2025), locomotion (Richards et al., 2025; Sharma et al., 2024), pain (J. Li et al., 2023; Singh et al., 2022; H. Wang et al., 2020), social behaviour (Y. Li et al., 2024), food seeking (Ye et al., 2023), and various aspects of fear and defensive behaviour (Z. Li et al., 2021; Lin et al., 2023; Richards et al., 2025; Venkataraman et al., 2019, 2021). The multifaceted roles and heterogeneous anatomical properties of the ZI suggest that the region may function as a sensory integration centre to produce appropriate behavioural output (X. Wang et al., 2019).

Numerous studies have begun to functionally characterise neurochemically defined ZI populations, however these generally fail to address the possibility of discrete functions across ZI sectors. Given that ZI sectors display unique hodological properties (Arena et al., 2024; Yang et al., 2022), it is likely that examining a neurochemically defined cell population in a sector indiscriminate manner may overlook nuances in ZI function. For example, a recent study demonstrated that two sector-specific subsets of GABAergic ZI neurons exhibited unique efferent projection patterns and different contributions to feeding and sleep/wake transitions (Zhu et al., 2025). Examining subpopulations of neurochemically defined ZI cells defined by sector location may further elucidate ZI function, and help reconcile various conflicting roles that have been attributed to this region.

We recently identified a genetically defined neuronal population expressing relaxin family peptide receptor 3 (RXFP3) spanning multiple ZI sectors and the adjacent lateral hypothalamus (LH; Richards et al., 2025). RXFP3 is a G_i/o_ protein-coupled receptor and the cognate receptor for relaxin-3 (Bathgate et al., 2006), a conserved neuropeptide predominantly synthesised in pontine nucleus incertus neurons (Burazin et al., 2002; S. Ma et al., 2007). We demonstrated that LH/ZI^RXFP3^ neurons project to multiple fear learning and defensive behaviour-implicated regions, notably, the lateral habenula (LHb), periaqueductal gray (PAG), and nucleus reuniens (Re). However, chemogenetically activating these cells during conditioned fear retrieval produced multiple behavioural phenotypes. This suggests that LH/ZI^RXFP3^ cells may consist of discrete subpopulations that mediate disparate behavioural responses to threats. In the current study, we demonstrate that topographically distinct subpopulations of LH/ZI^RXFP3^ neurons display unique brain-wide efferent connectivity patterns, especially to key regions involved in fear and defensive behaviour. These findings provide the first comprehensive neuroanatomical evidence for the existence of RXFP3+ subpopulations in the LH/ZI, providing a solid groundwork for future studies to parse out their functions.

## 2. Materials and Methods

### 2.1. Animals

Experiments were conducted in accordance with the Prevention of Cruelty to Animals Act (2004), under the guidelines of the National Health and Medical Research Council Code of Practice for the Care and Use of Animals for Experimental Purposes in Australia (8^th^ Edition, 2013) and approved by the Macquarie University Animal Ethics Committee (Animal Research Authority number: 2021/021). Inbred adult (8 – 13 weeks old) RXFP3-Cre mice (*n* = 21 female, 13 male; Ch’ng et al., 2019; Richards et al., 2025) were used in all experiments. Mice were group-housed (2-4 per cage) in individually ventilated chambers and maintained on a 12-hour light-dark cycle (lights on at 6 am) in a temperature-controlled environment (21°C ± 1°C) with nesting material and *ad libitum* access to standard chow and water. Seven mice (*n* = 6 female, 3 male) were excluded from final analyses due to misplaced viral injections.

### 2.2. Stereotaxic surgeries

Mice were anaesthetised under isoflurane (5% v/v in oxygen, maintained at 0.5% - 2%) and placed into a stereotaxic frame (David Kopf Instruments, CA, USA). Mice received a pre-operative injection of the non-steroidal anti-inflammatory analgesic carprofen (5 mg/kg, s.c., Rimadyl (Zoetis Australia)). For anterograde tracing experiments, mice were injected unilaterally with 25 nL (1 nL/sec) AAV-DJ-hSyn-FLEX-mGFP-synaptophysin-mRuby (diluted 1:10 in saline to 3 x 10^13^ GC/mL; obtained from Professor Andrew Allen, The University of Melbourne; Beier et al., 2015) at one of four different stereotaxic coordinates: the anterior LH (ALH; A/P: −1.00 mm, M/L: 1.25 mm, D/V: −5.05 mm), rostral ZI (ZIR; A/P: −0.95 mm, M/L: 0.75 mm, D/V: −4.55 mm), intermediate ZI (ZII; A/P: −1.60 mm, M/L: 1.00 mm, D/V: −4.50 mm), or caudal ZI (ZIC; A/P: −2.25 mm, M/L: 1.70 mm, D/V: −4.15 mm). For retrograde tracing experiments, mice were injected unilaterally with 40 nL (1 nL/sec) pENN.AAV.hSyn.HI.eGFP-Cre.WPRE-SV40 (2.1 x 10^13^ GC/mL; Addgene, 105540-AAVrg) into the ventrolateral periaqueductal gray (20° lateral angle, A/P: −4.30 mm, M/L: 0.35 mm, D/V: −2.85 mm) or the lateral habenula (20° lateral angle, A/P: −1.80 mm, M/L: 0.45 mm, D/V: −2.80 mm). During surgery, lidocaine was applied dropwise to the surgical site. Following infusion, the micropipette was left *in situ* (10 minutes), raised 0.1 mm, and left for a further 1 minute before removal. Injections were delivered with a Nanoject III Auto-Nanoliter Injector (3-000-207; Drummond Scientific Company, PA, USA).

### 2.3. Tissue preparation and histology

Two weeks following infusions to permit adequate viral transfection, all mice were anaesthetised with sodium pentobarbitone (80 mg/kg, i.p., Virbac, Australia). For anterograde tracing experiments, mice were transcardially perfused with heparinised saline at a flow rate of 7 ml/min for 2 minutes, followed by 5 minutes of 4% w/v paraformaldehyde (PFA) in 0.1 M phosphate-buffered saline (PBS). Brains were removed, post-fixed in 4% PFA in 0.1 M PBS (1 hr), washed with 0.1 M PBS (1 hr), and then placed in 30% w/v sucrose in 0.1 M PBS overnight for cryoprotection. Brains were snap-frozen over dry ice then sectioned coronally at 40 µm on a Leica CM1950 Cryostat (Leica Biosystems, Germany) and stored a 1-in-4 series in sodium azide (0.1% w/v in 0.1 M PBS) at 4 °C. For retrograde tracing experiments, mice were overdosed with sodium pentobarbitone (100 mg/kg, i.p., Virbac, Australia). Following euthanasia, brains were extracted and fresh frozen over dry ice. Brains were sectioned coronally at 8 µm on a Leica CM1950 Cryostat, slide-mounted onto Superfrost™ Plus slides (Epredia, NH, USA), and stored at −80 °C until required for RNAscope and fluorescent immunohistochemistry.

### 2.4. Immunohistochemistry (IHC)

For anterograde tracing experiments, fluorescent IHC was performed on every fourth section across the entire brain to amplify endogenous mGFP-labelled fibres and mRuby-labelled puncta, using procedures described previously (Walker et al., 2017) with appropriate modification to the antibodies used (Table 1).

**Table 1.** Antibodies used for anterograde tracing fluorescent immunohistochemistry.

| Antibody | Primary/<br>secondary | Dilution | Source | Cat. No. | RRID |
| --- | --- | --- | --- | --- | --- |
| Chicken anti-GFP polyclonal | Primary | 1:1000 | Abcam | Ab113 | AB_297905 |
| Rabbit anti-DsRed polyclonal | Primary | 1:1000 | Takara Bio<br>Clontech | 632496 | AB_10013483 |
| AF-488-conjugated anti-chicken IgG raised in donkey | Secondary | 1:500 | Jackson<br>ImmunoResearch | 703-545-155 | AB_2340375 |
| AF-555-conjugated anti-rabbit IgG raised in donkey | Secondary | 1:500 | Invitrogen | A32794 | AB_2762834 |

### 2.5. RNAscope^®^ fluorescent *in situ* hybridisation combined with fluorescent immunohistochemistry

For retrograde tracing experiments, RNAscope^®^ fluorescent *in situ* hybridisation was combined with fluorescent IHC to determine the precise origin and neurochemical phenotype of LH/ZI^RXFP3^ cells projecting to the lateral habenula and ventrolateral periaqueductal gray.

Slides were removed from −80 °C and sections were fixed in 4% PFA in PBS (15 minutes, RT) and washed twice in 0.1 M PBS (1 minute each, RT). After drying, a hydrophobic barrier was traced around each section before undergoing protease treatment (Protease Plus; 10 minutes, humid environment, 40 °C), then washed twice with dH_2_O. The RNAscope^®^ Multiplex Fluorescent V2 Assay (ACDBio, USA) was performed to label *Rxfp3* and S*lc17a6* mRNA (run 1) or *Rxfp3* and *Gad1* mRNA (run 2). All subsequent incubation steps were performed in a humid environment at 40 °C. Sections were rinsed twice with wash buffer (0.1x saline sodium citrate, 0.03% sodium dodecyl sulfate in dH_2_O) before probes for *Rxfp3* (Mm-*Rxfp3*-C2, #439381-C2; both runs), *Slc17a6* (Mm-*slc17a6*, #319171; run 1), and *Gad1* (Mm-*GAD1*-C3, #400951-C3; run 2) were applied and incubated for 90 minutes. A mouse-specific positive control probe (RNAscope^®^ 3-plex Positive Control Probe-Mm, #320881) and a universal negative control probe (RNAscope^®^ 3-plex Negative Control Probe, #320871) were applied to selected sections. Sections were then incubated in Amp1 (30 minutes), Amp2 (30 minutes), and Amp3 (15 minutes) to amplify target probes. Sections were rinsed in wash buffer after each Amp step (2x 2 minutes). For run 1, sections were then incubated in HRP C1 (15 minutes), Opal 690 (Akoya Biosciences, #FP1497001KT; 1:2000), and then HRP blocker (15 minutes) to develop the fluorophore signal for the *Slc17a6* probe. Sections were then incubated in HRP C2 (15 minutes), Opal 570 (Akoya Biosciences, #FP1488022KT; 1:1000), and then HRP blocker to develop the fluorophore signal for the *Rxfp3* probe. For run 2, sections were incubated in HRP C2, Opal 570 (1:1000), and HRP blocker to develop the *Rxfp3* signal first, then incubated in HRP C3, Opal 690 (1:2000), and HRP blocker to develop the fluorophore signal for the *Gad1* probe. For both runs, slides were then incubated in chicken anti-GFP polyclonal primary antibody (1:150; Abcam, ab113; RRID:AB_297905; 90 min, RT), washed twice in 0.1 M PBS (2x 2 min, RT), then incubated in AF-488-conjugated anti-chicken IgG raised in donkey secondary antibody (1:75; Jackson ImmunoResearch; 703-545-155, RRID: AB_2340375) for immunoamplification of the eGFP tag on the retrograde tracer virus. Sections were washed twice in 0.1 M PBS (2x 2 min, RT) and coverslipped with Fluoroshield™ with DAPI mounting medium (Sigma-Aldrich, MI, USA). Slides were left to dry overnight in the dark and stored at 4 °C until imaging. All RNAscope^®^ reagents were acquired from Advanced Cell Diagnostics, USA, unless otherwise indicated.

### 2.6. Microscopy and image acquisition

For anterograde tracing, overview images of sections were captured using an Olympus SLIDEVIEW™ VS200 Slide Scanner (Olympus/Evident, Japan; RRID:SCR_024783) with a UPLXAPO 10x/0.4 (WD = 3.1 mm) lens, Hamamatsu ORCA-Flash 4.0 CMOS camera, and VS200 ASW (v3.4.1; Olympus) imaging software. Alexa Fluor 488-labelled excitation was provided by a 475 nm LED from an X-Cite NOVEM light source (Excelitas, PA, USA). After reviewing the overview images in QuPath (v0.4.2) open-source software (Bankhead et al., 2017), brain regions of interest were chosen for high-magnification Z-stack imaging of both mGFP and mRuby immunofluorescence. Due to availability issues, two different confocal microscopes were used to image brain regions of interest. For cases 155, 156, 161, and 162, stitched Z-stacks were captured using an inverted Zeiss LSM 880 confocal microscope (Carl Zeiss AG, Germany) with a Plan-Apochromat 40x/1.3 NA oil objective using ZEN Black software. Photomicrographs were generated with 458 and 561 nm wavelength lasers to visualise Alexa Fluor 488- and 555-labelled signals, respectively. For all other cases, stitched Z-stacks were captured using a Leica Stellaris 5 confocal microscope (Leica Biosystems, Germany; RRID:SCR_024663) with a Plan-Apochromat 40x/1.3 NA oil objective, using Leica Application Suite (LAS) X software. Photomicrographs were generated with 499 and 553 nm wavelength lasers to visualise Alexa Fluor 488- and 555-labelled signals, respectively. A maximum intensity projection of each Z-stack was used for mGFP/mRuby density quantification and analysis.

For retrograde tracing, stitched confocal photomicrographs were captured using the Leica Stellaris 5 confocal microscope with a Plan-Apochromat 20x/0.75 NA objective using LAS X software. Photomicrographs were generated with 405, 499, 552, and 649 nm wavelength lasers to visualise DAPI-, Alexa Fluor 488-, Opal 570-, and Opal 690-labelled signals, respectively.

### 2.7. Mouse brain registration

For anterograde tracing, serial section overview images were registered to the Allen Mouse CCFv3 reference atlas (Q. Wang et al., 2020; RRID:SCR_020999) accounting for the angle of sectioning using QuickNII (v2.2; Puchades et al., 2019; RRID:SCR_016854). To account for tissue distortion, non-linear refinements were applied to registered slices using VisuAlign (v0.9; RRID:SCR_017978). Corresponding 40x high magnification Z-stacks were superimposed onto registered overview sections and regions with visible mGFP/mRuby expression were drawn onto 40x images in QuPath for subsequent mGFP and mRuby area quantification.

### 2.8. Quantification and analysis

For the anterograde tracing experiments, mGFP and mRuby area quantification was performed using QuPath’s in-built ‘pixel classifier’ function. A random subset of images from each experiment were assigned as training images for the ‘artificial neural network (ANN_MLP)’ pixel classifier, where areas of the training images were manually annotated as ‘positive’ or ‘negative’ for mGFP or mRuby expression until the machine learning algorithm produced an accurate profile of expression. Additional pixel classifiers were trained and applied to select regions of interest where the original classifiers were deemed inaccurate. Density measurements were calculated by taking the total detected area by the classifier and dividing it by the total area of each region of interest. For each brain region of interest, at least two measurements per mouse were included.

For the retrograde tracing experiments, LH and ZI regions were manually outlined and DAPI+ cells were batch detected by combining a custom script with the in-built ‘positive cell detection’ function. Detection of *Rxfp3*, *Slc17a6*, *Gad1* mRNA and eGFP immunoreactive cells were performed using QuPath’s in-built ‘object classifier’ function. Two images from each mouse were assigned as training images for the random trees object classifier, where cells were manually assigned as ‘positive’ or ‘negative’ for the marker of interest until the machine learning algorithm produced an accurate profile of expression. For *Rxfp3*, *Slc17a6*, and *Gad1*, a semi-quantitative method was employed, in which cells with two or more fluorescent dots within 5 µm of the DAPI-stained area were considered positive for the marker of interest (Ch’ng et al., 2019; Richards et al., 2025; Viden et al., 2022; Walker et al., 2021), with each dot denoting an individual mRNA molecule (F. Wang et al., 2012). Trained classifiers were batch-applied to all outlined regions for all images using a custom script, which classified each DAPI+ cell as being ‘positive’ or ‘negative’ for each marker. QuPath scripts can be found at https://github.com/BrandonKR1.

### 2.9. Neuroanatomical nomenclature

The abbreviations in Table 2 follow those in the Mouse Brain Atlas in Stereotaxic Coordinates (Paxinos & Franklin, 2004) or the Allen Mouse CCFv3 Reference Atlas (Q. Wang et al., 2020), with some exceptions. The bed nucleus of the stria terminalis was subdivided into a ventral aspect (BSTV) and a caudal aspect (BSTC), given the lack of mGFP and mRuby immunoreactivity observed in the dorsal and rostral parts of the structure. The lateral hypothalamic area was divided into three rostrocaudal zones (anterior, tuberal, and mammillary) according to the nomenclature of Hahn et al. (2019). The rostral periaqueductal gray (RPAG) was used to delineate the PAG before the columnar organisation of the nucleus became apparent.

**Table 2.** List of abbreviations.

|  |  |
| --- | --- |
| **abducens nucleus** | **6N** |
| **accumbens nucleus** | Acb |
| **anterior hypothalamic area** | AH |
| **anterior part of the lateral hypothalamic area** | ALH |
| **anteromedial thalamic nucleus, ventral part** | AMV |
| **anterior pretectal nucleus** | APT |
| **bed nucleus of the stria terminalis, caudal part** | BSTC |
| **bed nucleus of the stria terminalis, ventral part** | BSTV |
| **central gray of the pons** | CGPn |
| **centrolateral thalamic nucleus** | CL |
| **central medial thalamic nucleus** | CM |
| **cuneiform nucleus** | CnF |
| **nucleus of Darkschewitsch** | Dk |
| **dorsal lateral geniculate nucleus** | DLG |
| **dorsolateral periaqueductal gray** | DLPAG |
| **dorsomedial hypothalamic nucleus** | DM |
| **dorsomedial part of the lateral hypothalamic area** | dmLH |
| **dorsomedial periaqueductal gray** | DMPAG |
| **deep gray layer of the superior colliculus** | DpG |
| **dorsal paragigantocellular nucleus** | DPGi |
| **dorsal raphe nucleus** | DR |
| **dorsal tegmental nucleus** | DTg |
| **ethmoid thalamic nucleus** | Eth |
| **Edinger-Westphal nucleus** | EW |
| **fields of Forel** | FF |
| **fasciculus retroflexus** | fr |
| **gigantocellular reticular nucleus** | Gi |
| **intergeniculate leaf** | IGL |
| **incertohypothalamic area** | IHy |
| **intermediate part of the lateral hypothalamic area** | iLH |
| **intermediodorsal thalamic nucleus** | IMD |
| **interstitial nucleus of Cajal** | InC |
| **intermediate gray layer of the superior colliculus** | InG |
| **intermediate white layer of the superior colliculus** | InWh |
| **inferior olive** | IO |
| **intermediate reticular nucleus** | IRt |
| **laterodorsal thalamic nucleus** | LD |
| **laterodorsal tegmental nucleus** | LDTg |
| **lateral habenula** | **LHb** |
| **lateral division of the lateral habenula** | **LHbL** |
| **oval subnucleus of the lateral division of the lateral habenula** | **LHbLO** |
| **medial division of the lateral habenula** | **LHbM** |
| **parvocellular subnucleus of the medial division of the lateral habenula** | **LHbMPc** |
| **lateral posterior thalamic nucleus** | **LP** |
| **lateral periaqueductal gray** | **LPAG** |
| **lateral paragigantocellular nucleus** | **LPGi** |
| **lateral preoptic area** | **LPO** |
| **lateral septal nucleus** | **LS** |
| **magnocellular reticular nucleus** | **MARN** |
| **mediodorsal thalamic nucleus** | **MD** |
| **medial geniculate nucleus** | **MG** |
| **medial lemniscus** | **ml** |
| **median raphe nucleus** | **MnR** |
| **medial preoptic area** | **MPO** |
| **medial pretectal nucleus** | **MPT** |
| **midbrain reticular nucleus** | **MRN** |
| **medial septal nucleus** | **MS** |
| **nucleus of the diagonal band** | **NDB** |
| **nucleus incertus** | **NI** |
| **nucleus of the lateral lemniscus** | **NLL** |
| **optic nerve layer of the superior colliculus** | **Op** |
| **olivary pretectal nucleus** | **OPT** |
| **periaqueductal gray** | **PAG** |
| **phosphate-buffered saline** | **PBS** |
| **paracentral thalamic nucleus** | **PC** |
| **nucleus of the posterior commissure** | **PCom** |
| **posterodorsal tegmental nucleus** | **PDTg** |
| **parafascicular thalamic nucleus** | **PF** |
| **paraformaldehyde** | **PFA** |
| **posterior hypothalamic area** | **PH** |
| **posterior part of the lateral hypothalamic area** | **PLH** |
| **preammillary nucleus, dorsal part** | **PMD** |
| **pontine reticular nucleus caudal part** | **PnC** |
| **pontine reticular nucleus, oral part** | **PnO** |
| **posterior thalamic nuclear group** | **Po** |
| **posterior thalamic nuclear group, triangular part** | **PoT** |
| **posterior pretectal nucleus** | **PPT** |
| **pedunculo-pontine tegmental nucleus** | **PPTg** |
| **parapyramidal nucleus** | **PPy** |
| **prepositus nucleus** | **Pr** |
| **precommissural nucleus** | **PrC** |
| **parastrial nucleus** | **PS** |
| **parasubthalamic nucleus** | **PSTh** |
| **paratenial thalamic nucleus** | **PT** |
| **paraventricular thalamic nucleus** | **PV** |
| **paraventricular hypothalamus** | **PVH** |
| **pyramidal tract** | **py** |
| **red nucleus** | **R** |
| **reuniens thalamic nucleus** | **Re** |
| **raphe magnus nucleus** | **RMg** |
| **rostral periaqueductal gray** | **RPAG** |
| **retroparafascicular nucleus** | **RPF** |
| **nucleus raphe pontis** | **RPO** |
| **room temperature** | **RT** |
| **reticular thalamic nucleus** | **Rt** |
| **reticulotegmental nucleus of the pons** | **RtTg** |
| **sagulum nucleus** | **Sag** |
| **superior cerebellar peduncle** | **scp** |
| **substantia innominata** | **SI** |
| **sublaterodorsal nucleus** | **SLD** |
| **stria medullaris of the thalamus** | **sm** |
| **substantia nigra, compact part** | **SNC** |
| **subparafascicular thalamic nucleus** | **SPF** |
| **subthalamic nucleus** | **STh** |
| **supraoculomotor periaqueductal gray** | **Su3** |
| **subcoeruleus nucleus** | **SubC** |
| **superficial gray layer of the superior colliculus** | **SuG** |
| **supramammillary nucleus** | **SuM** |
| **trapezoid body** | **tz** |
| **ventral anterior thalamic nucleus** | **VA** |
| **ventrolateral thalamic nucleus** | **VL** |
| **ventral lateral geniculate nucleus** | **VLG** |
| **ventrolateral periaqueductal gray** | **VLPAG** |
| **ventromedial thalamic nucleus** | **VM** |
| **ventromedial hypothalamic nucleus** | **VMH** |
| **ventral posterolateral thalamic nucleus** | **VPL** |
| **ventral posteromedial thalamic nucleus** | **VPM** |
| **ventral tegmental area** | **VTA** |
| **ventral tegmental nucleus** | **VTg** |
| **zona incerta, caudal** | **ZIC** |
| **zona incerta, dorsal/ventral** | **ZID/ZIV** |
| **zona incerta, intermediate** | **ZII** |
| **zona incerta, rostral** | **ZIR** |

## 3. Results

### 3.1. Injection sites

To determine whether subpopulations of LH/ZI^RXFP3^ cells exhibit distinct brain-wide projection patterns, we unilaterally targeted a Cre-dependent anterograde tracer (AAV-DJ-hSyn-FLEX-mGFP-synaptophysin-mRuby; Figure 1A) to four different areas of the LH/ZI in RXFP3-Cre mice: the anterior lateral hypothalamic area (ALH^RXFP3^; *n* = 3), rostral zona incerta (ZIR^RXFP3^; *n* = 4), intermediate zona incerta (ZII^RXFP3^; *n* = 6), and caudal zona incerta (ZIC^RXFP3^; *n* = 4). Although both males and females were used, sex differences were not analysed due to unequal distribution of sex across groups (Supplementary Table 1).

**Figure 1.**
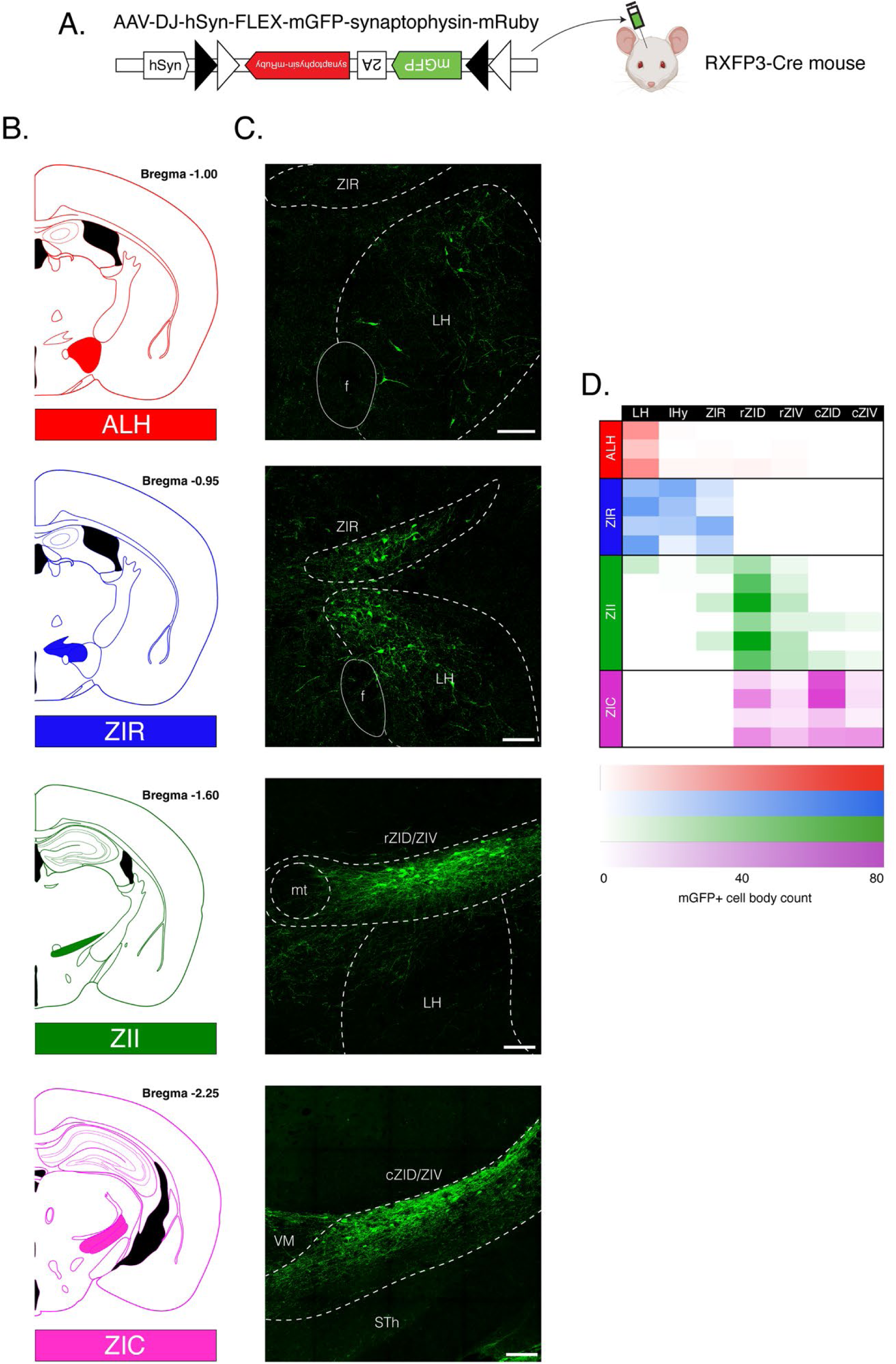
Overview of injection sites for whole-brain anterograde tracing of RXFP3+ cells from topographically distinct areas of the LH/ZI. (A) Schematic of the anterograde tracing strategy. A Cre-dependent anterograde tracer virus was injected into RXFP3-Cre mice to trace the efferent projections of RXFP3 cells. In transduced cells, mGFP expression is observed in the cell body and efferent fibres, while mRuby expression is driven by the presence of synaptophysin at pre-synaptic terminals. (B) Schematics depicting distinct areas of the LH/ZI targeted with the anterograde tracer: anterior lateral hypothalamic area (ALH; red), rostral zona incerta (ZIR; blue), intermediate zona incerta (ZII; green), caudal zona incerta (ZIC; pink). (C) Representative confocal photomicrographs of mGFP expression at the focal injection site. (D) Heat map showing the distribution of mGFP+ cell bodies colour-coded by group. Each row represents one mouse. cZID, caudal part of the dorsal zona incerta; cZIV, caudal part of the ventral zona incerta; f, fornix; IHy, incertohypothalamic area; LH, lateral hypothalamus; mt, mammillothalamic tract; rZID, rostral part of the dorsal zona incerta; rZIV, rostral part of the ventral zona incerta; STh, subthalamic nucleus; VM, ventromedial thalamic nucleus; ZIR, zona incerta, rostral. *n* = 3-6/group. Scale bars = 100 µm.

In ALH targeted injections, mGFP+ cells sparsely populated the ALH and were largely restricted to this area (Figure 1B – D, top row, Supplementary Figure 1). In ZIR targeted injections, a dense, contiguous group of mGFP+ cells spanning the ZIR and dorsomedial part of the lateral hypothalamic area (dmLH) was observed (Figure 1B – D, second row, Supplementary Figure 2). In ZIR^RXFP3^ cases, mGFP+ cells were observed between the ZIR and dmLH immediately dorsolateral to the fornix (Supplementary Figure 2). For clarity, we have defined this zone as the incertohypothalamic area (IHy; Figure 1D), as it is undefined in common mouse brain atlases (Paxinos & Franklin, 2004; Q. Wang et al., 2020). In ZII targeted injections, transduced mGFP+ cells primarily populated the rostral half of the ZID/ZIV, but were occasionally observed near the ZIR-ZID/ZIV border (Figure 1D, third row, Supplementary Figure 3). mGFP+ cells were more numerous in the rostral half of the ZID (62.5% ± 5.3% of total mGFP+ cells) than in the rostral half of the ZIV (17.3% ± 2.1%).

Similarly, in ZIC targeted injections, mGFP+ labelled cells predominated in the ZID (72.1% ± 6.4%) than in the ZIV (27.9% ± 6.4%). However, the caudal end of the rostral ZID/ZIV also contained mGFP+ cells (38.8% ± 5.1%; Figure 1B – D, bottom row; Supplementary Figure 4).

### 3.2. Macroscale efferent connectivity patterns

We first examined whether LH/ZI^RXFP3^ cells exhibited distinct macroscale efferent connectivity patterns by analysing the percentage of mGFP+ fibres and mRuby+ boutons in each major brain subdivision relative to brain-wide mGFP+/mRuby+ expression for each group. Of these two measures, mRuby+ immunoreactivity provides the more faithful indicator of monosynaptic connections, since mGFP+ immunoreactivity could also indicate a fibre that passes through but does not terminate in that region. Therefore, for brevity we will restrict this description to mRuby+ values, however both measures can be seen in Figure 2.

**Figure 2.**
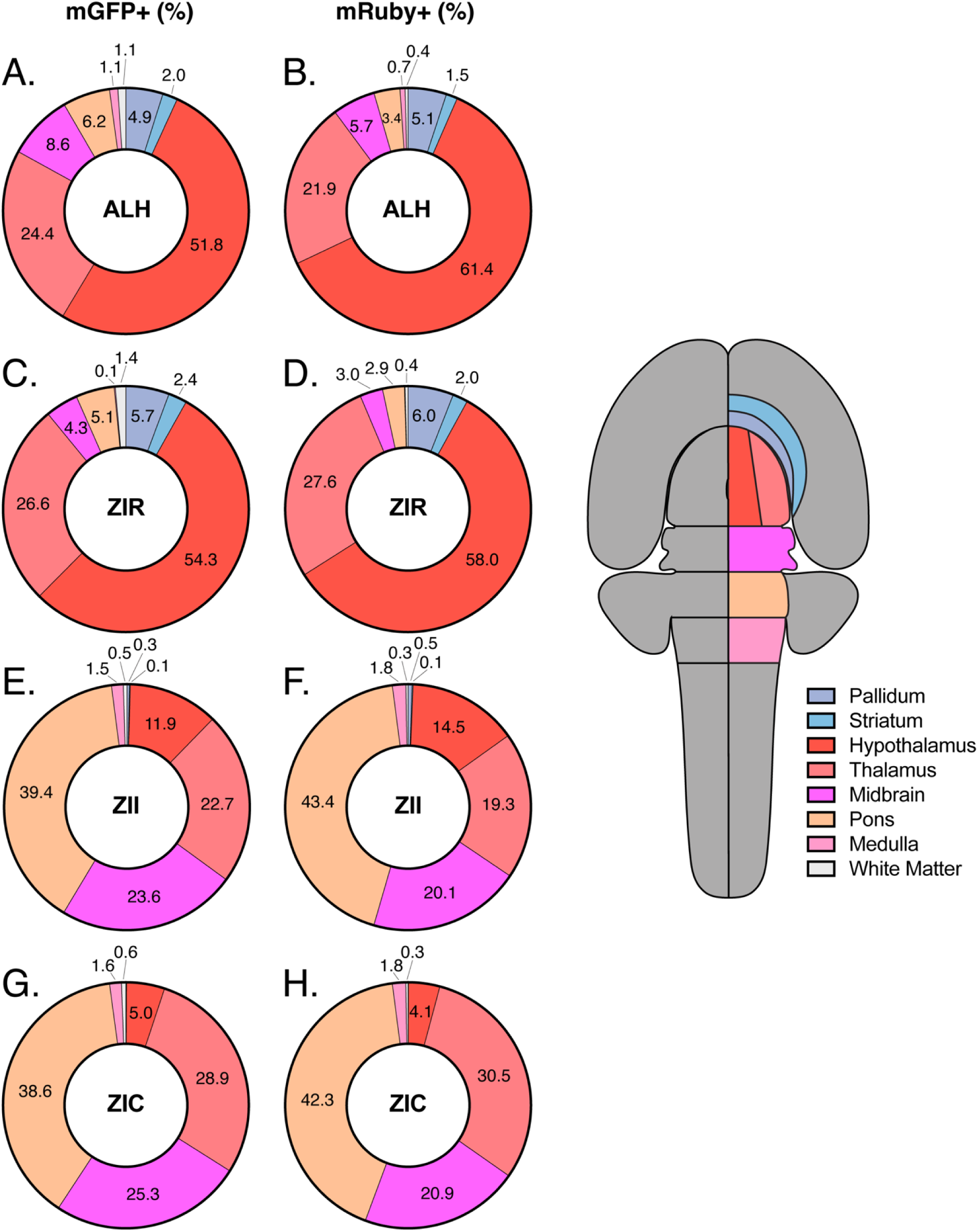
Topographically distinct LH/ZI^RXFP3^ cells display unique efferent projection patterns to major subdivisions of the brain. Donut graphs depict the average percentage of mGFP+ fibres (A, C, E, G) and mRuby+ boutons (B, D, F, H) observed in each major brain subdivision for ALH^RXFP3^ cases (A, B), ZIR^RXFP3^ cases (C, D), ZII^RXFP3^ cases (E, F), and ZIC^RXFP3^ cases as a proportion of overall mGFP+ and mRuby+ expression. Donut graph segment colours represent the major brain subdivisions shown in the flatmap (right). *n* = 3-6/group. Individual case data can be found in Supplementary Table 2.

ALH^RXFP3^ and ZIR^RXFP3^ cases primarily innervated diencephalic regions (ALH^RXFP3^: 83.3% ± 1.9%; ZIR^RXFP3^: 85.6% ± 1.7%) and showed modest projections to pallidal and striatal regions of the forebrain (ALH^RXFP3^: 6.6% ± 2.8%; ZIR^RXFP3^: 8.0% ± 2.5%). Conversely, ZII^RXFP3^ and ZIC^RXFP3^ cases largely avoided these areas (ZII^RXFP3^: 0.6% ± 0.3%; ZIC^RXFP3^: 0.1% ± 0.01%) and mainly projected to the midbrain and hindbrain (ZII^RXFP3^: 65.3% ± 10.2%; ZIC^RXFP3^: 65.0% ± 9.8%), with robust input to the pons (ZII^RXFP3^: 43.4% ± 7.3%; ZIC^RXFP3^: 42.3% ± 7.8%). Although there were similar proportions of intra-diencephalic projections between ZII^RXFP3^ and ZIC^RXFP3^ cases (ZII^RXFP3^: 33.8% ± 10.0%; ZIC^RXFP3^: 34.6% ± 9.9%), ZIC^RXFP3^ cases showed a bias towards innervating the thalamus rather than the hypothalamus (30.5% ± 10.7% thalamic input; 4.1% ± 1.0% hypothalamic input), whereas ZII^RXFP3^ cases displayed more balance (19.3% ± 5.1% thalamic input; 14.5% ± 5.4% hypothalamic input). In all cases, no projections were observed in the hippocampus, amygdala, and cerebral cortex. Cerebellar nuclei were not captured during tissue processing.

### 3.3. Brain-wide distribution of mGFP+ fibres and mRuby+ boutons

To quantify mGFP+ fibre and mRuby+ bouton density throughout the brain, we calculated the average mGFP+/mRuby+ area as a proportion of the total area for each brain region analysed, for each case (Supplementary Figure 5, 6), then averaged these values across the injection site group (Figure 3). We then categorised these density values into a 7-point ordinal scale to produce density heatmaps (Figure 3, Supplementary Figure 5, 6).

**Figure 3.**
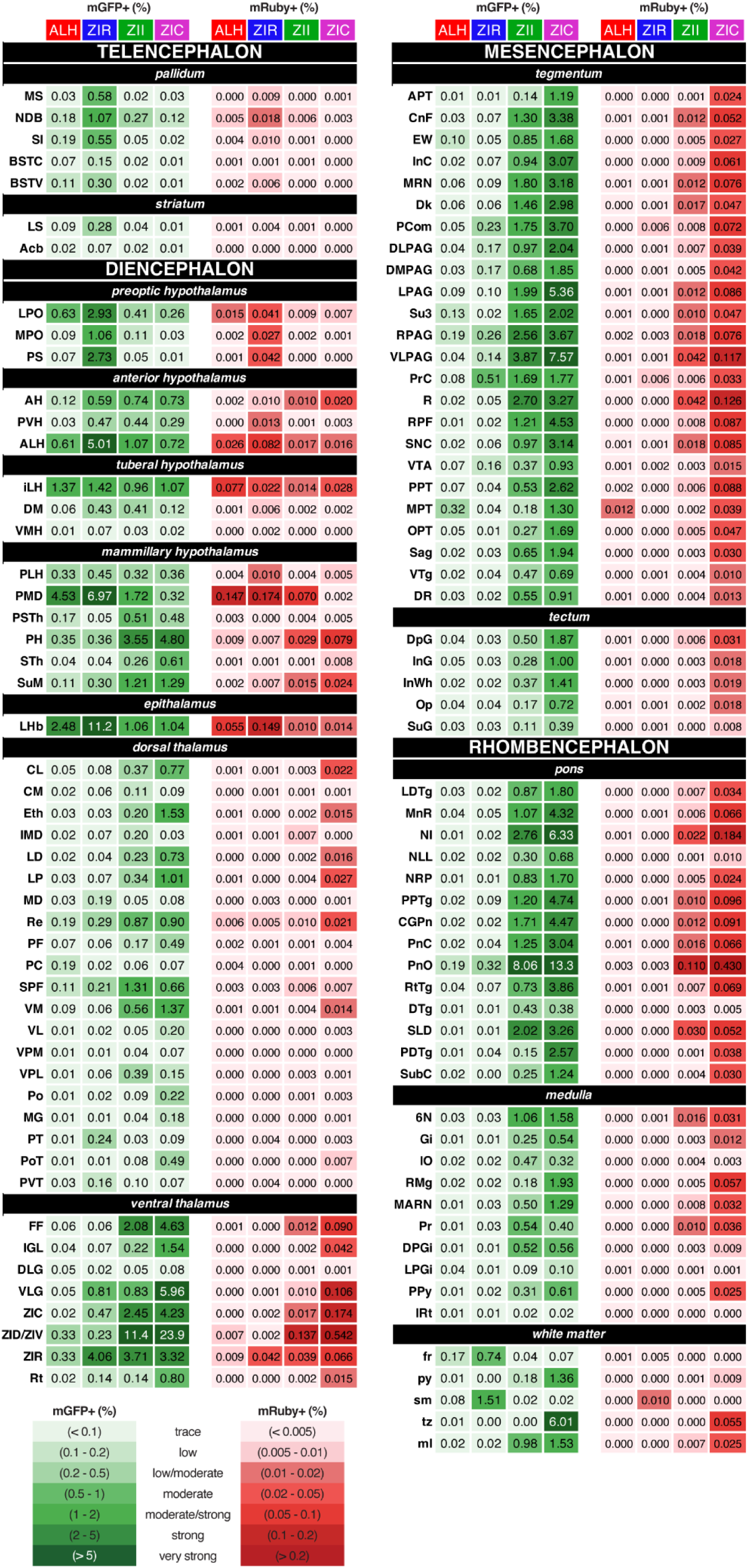
Heatmap of topographically distinct LH/ZI^RXFP3^ efferent projections. Green columns represent mGFP+ fibre density, whereas red columns represent mRuby+ bouton density. Numbers are the average mGFP+/mRuby+ area expressed as a proportion of the total area averaged across the injection site group.

The following descriptions refer to the mGFP+ fibre patterns, since these provide more descriptive data. Typically, mRuby+ expression mirrored mGFP+ expression but was less dense; any discrepancy from this pattern is mentioned in the text. Only ipsilateral projections were quantified; if substantial contralateral projections were observed, these were also noted.

#### 3.3.1. Telencephalon

##### 3.3.1.1. Pallidum

Most efferents to the pallidum were observed in two out of four ZIR^RXFP3^ cases (#164, #165). The strongest projections were to the nucleus of the diagonal band (NDB; Figure 4). Here, diagonally oriented fibres occupied the rostrocaudal extent of the NDB but were more densely packed in the intermediate areas of the nucleus (∼Bregma +0.60). Moderate density projections to the medial septal nucleus (MS) and substantia innominata (SI) were also observed in these cases (Figure 4), while low-density projections to the bed nucleus of the stria terminalis, ventral part (BSTV), were observed across all ZIR^RXFP3^ cases. In most ALH^RXFP3^, ZII^RXFP3^, and ZIC^RXFP3^ cases, labelling was sparse or low in the analysed pallidal nuclei.

**Figure 4.**
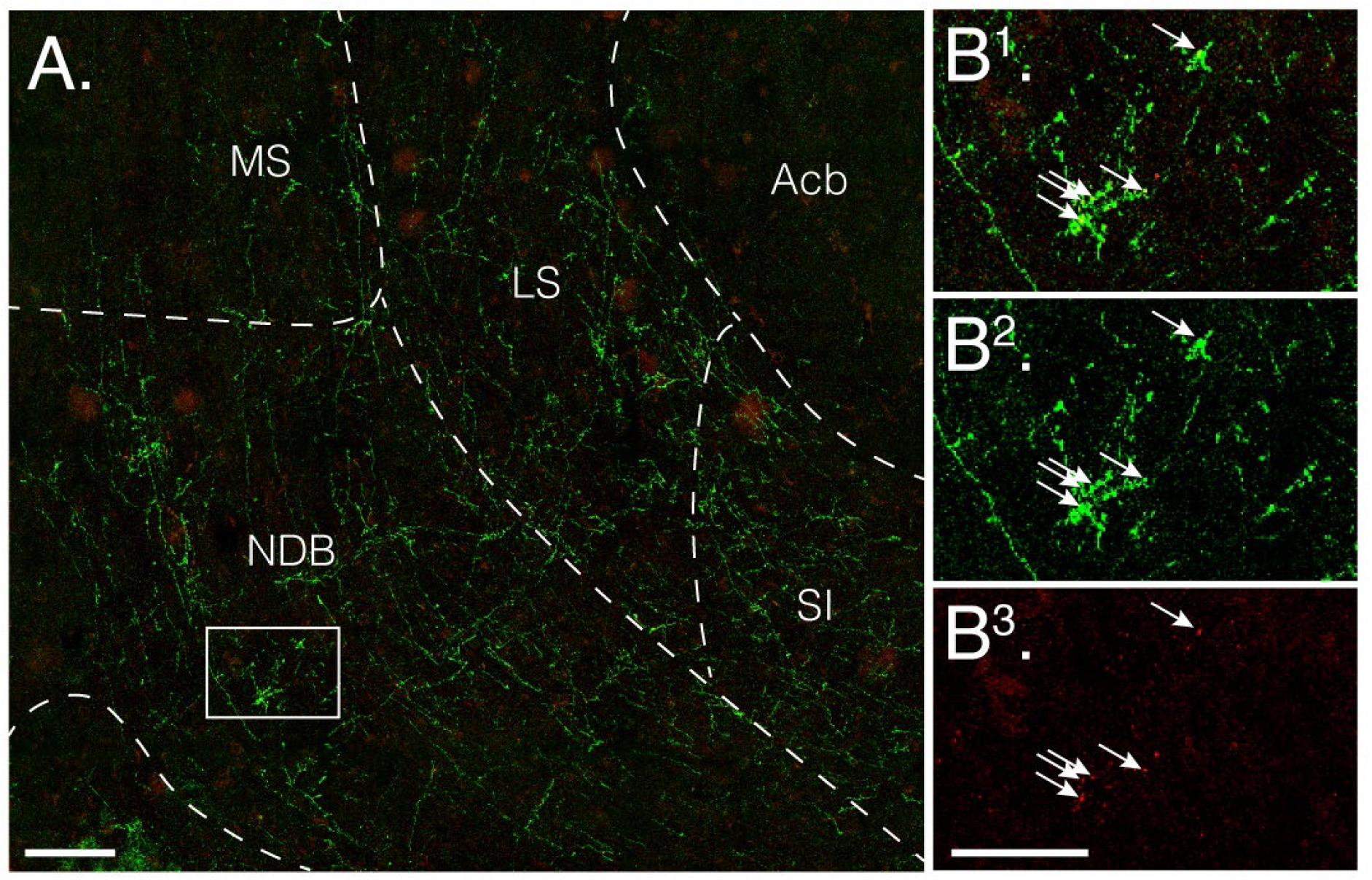
ZIR^RXFP3^ cases project to the ventral telencephalon. (A) Stitched fluorescent confocal photomicrograph of mGFP+ immunoreactive fibres (green) and mRuby+ immunoreactive boutons (red) in the medial septal nucleus (MS), ventral part of the lateral septal nucleus (LS), nucleus of the diagonal band (NDB), and medial part of the substantia innominata (SI) from ZIR^RXFP3^ case #165. Single-channel fluorescent confocal photomicrographs from the inset box in A are shown in panel B^1^ (merge), B^2^ (mGFP), and B^3^ (mRuby), showing clusters of pre-synaptic terminals in the DB; white arrows indicate some examples. Scale bars: 100 µm (A); 50 µm (B). Acb, accumbens nucleus.

##### 3.3.1.2. Striatum

Independent of group, labelling was generally absent/sparse throughout the striatum. However, one ZIR^RXFP3^ case (#165) produced moderate density labelling confined to the ventral third of the lateral septal nucleus (LS; Figure 4) and low/moderate labelling along the medial border of the accumbens nucleus (Acb).

#### 3.3.2. Diencephalon

##### 3.3.2.1. Preoptic hypothalamus

Projections to the preoptic hypothalamus were most prominent in ZIR^RXFP3^ cases. Here, a dense network of fibres blanketed most of the lateral preoptic area (LPO), continuous with labelling in the lateral part of the medial preoptic area (MPO), the parastrial nucleus (PS), and the lateral SI (Figure 5A). In ALH^RXFP3^, ZII^RXFP3^, and ZIC^RXFP3^ cases, low/moderate to moderate labelling occupied the rostrocaudal extent of the LPO, especially on its ventromedial side (Figure 5B), while sparse projections were observed throughout the rest of the preoptic hypothalamus.

**Figure 5.**
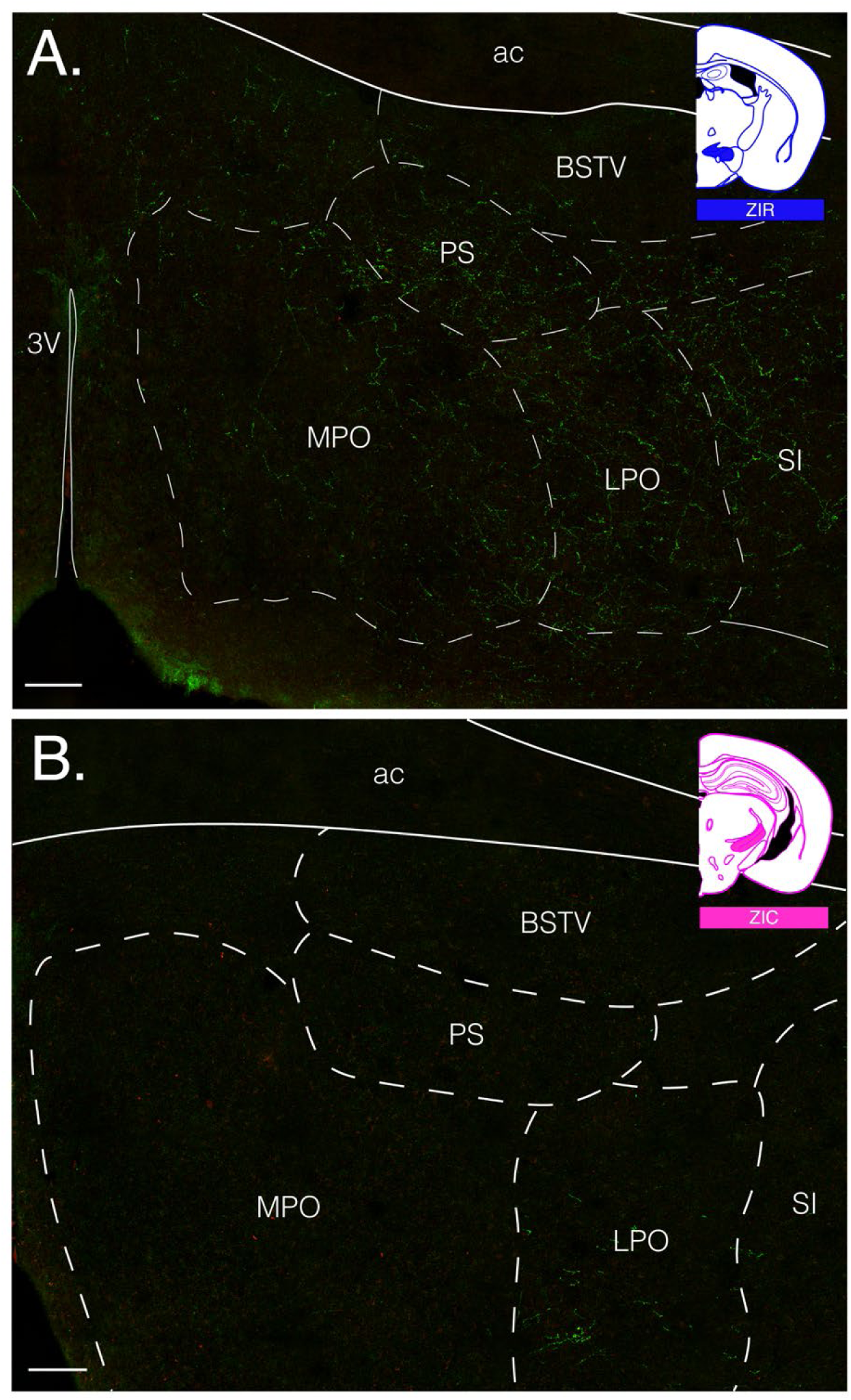
ZIR^RXFP3^ and ZIC^RXFP3^ cases display different projection patterns to the preoptic hypothalamus. Stitched confocal photomicrographs of mGFP+ immunoreactive fibres (green) and mRuby+ immunoreactive boutons (red) in the preoptic hypothalamus from a ZIR^RXFP3^ case (A) and a ZIC^RXFP3^ case (B). 3V, third ventricle; ac, anterior commissure; BSTV, bed nucleus of the stria terminalis, ventral part; MPO, medial preoptic area; LPO, lateral preoptic area; PS, parastrial nucleus; SI, substantia innominata. Scale bars = 100 µm.

##### 3.3.2.2. Anterior hypothalamus

The anterior hypothalamus generally received moderate input and displayed a consistent pattern of labelling independent of group. Fibres often ran diagonally and formed a continuous pathway spanning the medial aspect of the ALH and the lateral aspect of the anterior hypothalamic area (AH), with some fibres passing through the fornix (Figure 6A). Rostrally, labelling was absent in the paraventricular hypothalamus (PVH). Caudally (∼Bregma −1.2), a band of fibres traversed the dorsal border of the AH without invading the suprajacent PVH (Figure 6A). Though low/moderate mGFP+ labelling was observed in the PVH caudally (except for ALH^RXFP3^ cases), mRuby+ immunoreactivity was generally absent/sparse in the PVH, indicating that these fibres were likely *en passant*. In most ZII^RXFP3^ cases, a very dense network of fibres continuous with the ZIR populated the undifferentiated zone between the ZIR and dmLH, consistent with the incertohypothalamic area (IHy) as described by Sita and colleagues (2007; Figure 6B). ZIR^RXFP3^ cases produced very strong densities of quantified mGFP+ immunoreactivity in the ALH, mainly because of the mGFP+ cell bodies marking the injection site.

**Figure 6.**
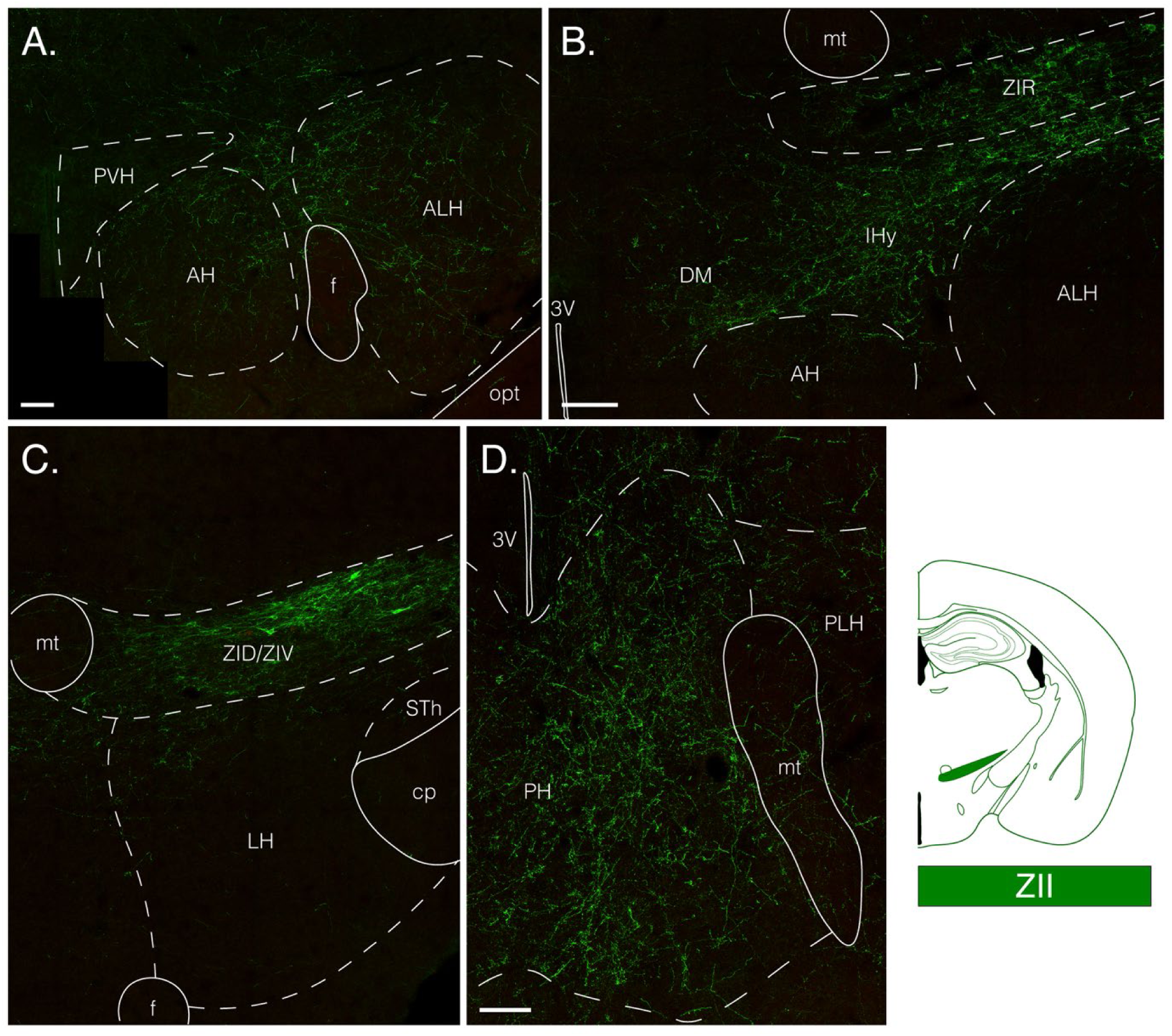
ZII^RXFP3^ projections to the hypothalamus. Representative stitched fluorescent confocal photomicrographs of mGFP+ fibres (green) and mRuby+ boutons (red) in the anterior hypothalamus (A), the anterior/tuberal hypothalamus border (B), tuberal hypothalamus (C), and the rostral part of the mammillary hypothalamus (D) for ZII^RXFP3^ cases. 3V, third ventricle; AH, anterior hypothalamic area; ALH, anterior part of the lateral hypothalamic area; cp, cerebral peduncle; DMH, dorsomedial hypothalamic nucleus; f, fornix; IHy, incertohypothalamic area; LH, lateral hypothalamic area; mt, mammillothalamic tract; opt, optic tract; PH, posterior hypothalamic area; PLH, posterior part of the lateral hypothalamic area; PVH, paraventricular hypothalamic nucleus; STh, subthalamic nucleus; ZID/ZIV, zona incerta, dorsal/ventral part; zona incerta, rostral part. All scale bars = 100 µm.

##### 3.3.2.3. Tuberal hypothalamus

The tuberal hypothalamus showed similar labelling patterns across groups. Sparse, mostly non-overlapping fibres spanned the intermediate part of the lateral hypothalamic area (iLH). However, in ZII^RXFP3^ cases, a dense plexus additionally occupied the dorsal iLH, continuous with ZIV expression (Figure 6C). Only sparse expression was observed in the ventromedial hypothalamic nucleus (VMH) across all groups. In ZIR^RXFP3^ and ZII^RXFP3^ cases, moderate density fibres were observed in the rostral part of the dorsomedial hypothalamic nucleus (DM), continuing medially from the ZI and IHy (Figure 6B).

##### 3.3.2.4. Mammillary hypothalamus

Within the mammillary hypothalamus, there were several sub-region differences in efferent projection patterns between injection sites. The most marked difference was observed in the premammillary nucleus, dorsal part (PMD), where robust labelling occupied most of the region in both ALH^RXFP3^ and ZIR^RXFP3^ cases (Figure 7A, B). In contrast, only low-density labelling was observed there in ZIC^RXFP3^ cases (Figure 7E). Low-density labelling was also observed in the PMD in ZII^RXFP3^ cases (Figure 7C), except for case #179, which exhibited strong labelling analogous to ALH^RXFP3^ and ZIR^RXFP3^ cases (Figure 7D). In all cases, slightly weaker labelling was found in the contralateral PMD. As case #179 was the only ZII^RXFP3^ case with some transfected cell bodies in the LH, the source of observed PMD efferents likely arose from LH^RXFP3^ cells.

**Figure 7.**
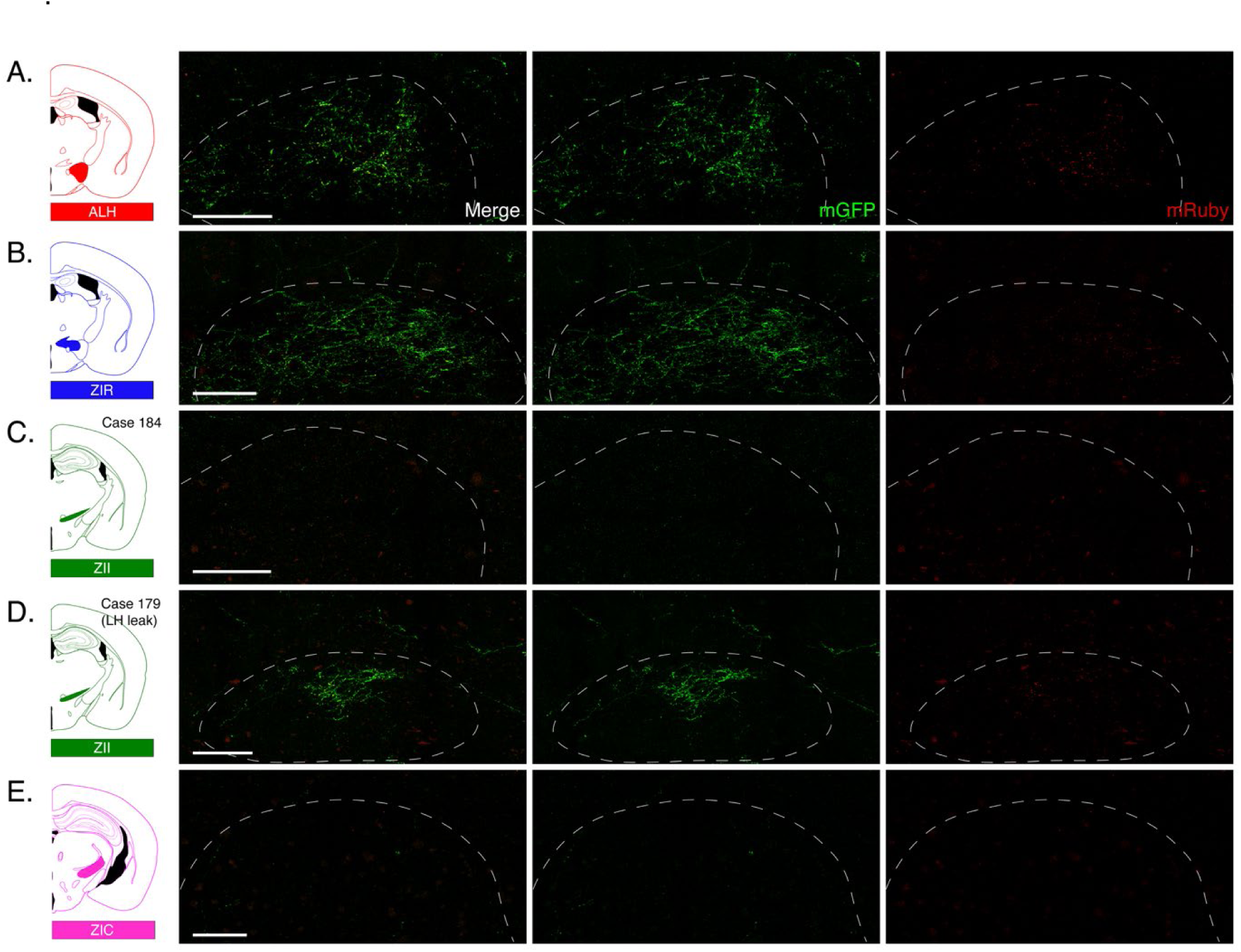
LH/ZI^RXFP3^ cells display distinct projection patterns to the premammillary nucleus, dorsal part (PMD). Representative stitched fluorescent confocal photomicrographs of mGFP+/mRuby+ expression (left), mGFP+ only (middle), and mRuby+ only (right) in the PMD in an ALH^RXFP3^ case (A), ZIR^RXFP3^ case (B), ZII^RXFP3^ case (C), a ZII case with some transfected cell bodies in the LH (D), and a ZIC^RXFP3^ case (E). All scale bars = 100 µm.

Intense labelling was observed in the posterior hypothalamic area (PH) in ZII^RXFP3^ (Figure 6D) and ZIC^RXFP3^ cases, while only low/moderate labelling was observed in ALH^RXFP3^ and ZIR^RXFP3^ cases. Rostrally, expression was biased towards the dorsomedial PH, but occupied most of the nucleus caudally. Labelling in the supramammillary nucleus (SuM) was mainly restricted to its medial area and was continuous with PH expression. Both the subthalamic nucleus (STh) and parasubthalamic nucleus (PSTh) received low/moderate input from some ZII^RXFP3^ and ZIC^RXFP3^ cases but did not receive input from ZIR^RXFP3^ or ALH^RXFP3^ cases.

##### 3.3.2.5. Lateral habenula

The epithalamic lateral habenula (LHb) was a key ipsilateral and contralateral target, particularly in ZIR^RXFP3^ and ALH^RXFP3^ cases. ZIR^RXFP3^ cases displayed the highest density of mGFP+ and mRuby+ immunoreactivity across all groups, especially at the caudal end (Figure 8A). Inputs to the LHb from ZIR^RXFP3^ and ALH^RXFP3^ cases comprised large proportions of the total mGFP+ and mRuby+ area across the brain. Indeed, the LHb accounted for ∼15% of the total mGFP+ area (more than half of the overall thalamic input) and ∼21% of the total mRuby+ area (more than three-quarters of the overall thalamic input) in ZIR^RXFP3^ cases (Figure 8B). The LHb exhibited ∼13% of the brain-wide total mGFP+ area (about half of the overall thalamic input) and ∼12% of the brain-wide total mRuby+ area (more than half of the overall thalamic input) in ALH^RXFP3^ cases.

**Figure 8.**
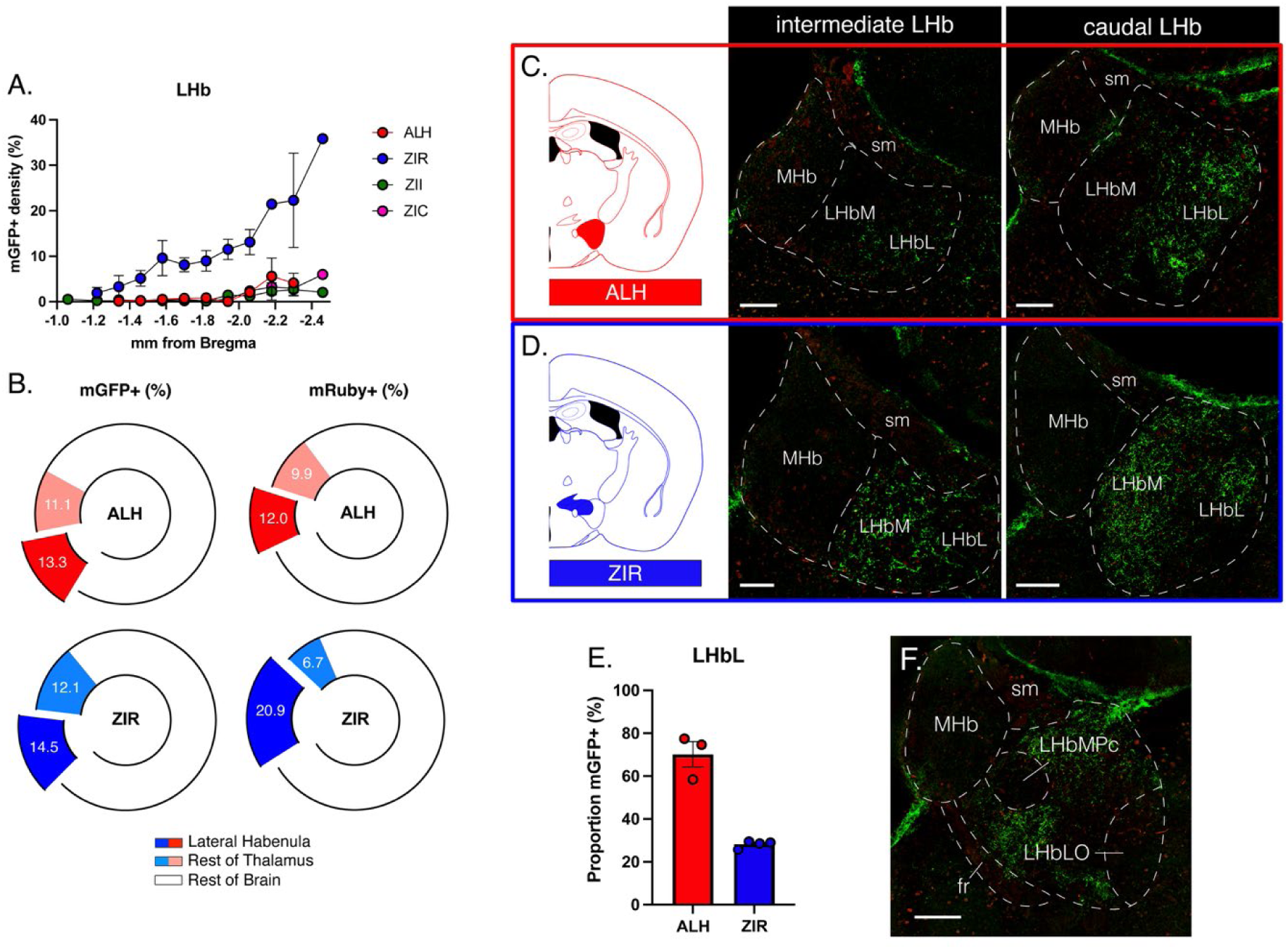
ALH^RXFP3^ cells and ZIR^RXFP3^ cases display distinct, strong projections to the lateral habenula (LHb). (A) Graph showing the average mGFP+ density divided by Bregma level for each injection site group. (B) Donut graphs showing the average relative percentage of mGFP+ fibres (left) and mRuby+ boutons (right) observed in the LHb (dark red, dark blue) and the rest of the thalamus (light red, light blue) in both ALH^RXFP3^ cases (red) and ZIR^RXFP3^ cases as a proportion of total observed mGFP+/mRuby observed throughout the entire brain. (C, D) Representative stitched confocal photomicrographs of mGFP/mRuby expression in the intermediate part of the LHb (left) and caudal part of the LHb (right) in ALH^RXFP3^ cases (C) and ZIR^RXFP3^ cases. (E) Bar graph indicating the average proportion of mGFP+ expression in the lateral part of the LHb for both ALH^RXFP3^ cases (red) and ZIR^RXFP3^ cases (blue). (F) Representative stitched confocal photomicrograph of the caudal LHb of a ZIR^RXFP3^ case demonstrating the subnuclear organisation of the LHb. Data are presented as mean ± SEM. Scale bars = 100 µm. fr, fasciculus retroflexus; MHb, medial habenula; LHbL, lateral division of the lateral habenula; LHbLO, oval subnucleus of the lateral division of the lateral habenula; LHbM, medial division of the lateral habenula; LHbMPc, parvocellular subnucleus of the medial division of the lateral habenula; sm, stria medullaris of the thalamus.

Although ALH^RXFP3^ and ZIR^RXFP3^ cases exhibited similar densities of mGFP/mRuby expression, they displayed unique innervation patterns. Both showed moderate labelling in the rostral LHb, which was not circumscribed to a particular subregion of the nucleus.

However, in the intermediate and caudal LHb, ZIR^RXFP3^ cases strongly innervated the medial half of the LHb (LHbM), whereas ALH^RXFP3^ cases mostly innervated the lateral half of the LHb (LHbL; Figure 8C, D). Indeed, only 27.4% (± 0.9%) of the observed LHb mGFP+ immunoreactivity occupied the LHbL in ZIR^RXFP3^ cases compared to 70.1% (± 5.9%) of the LHbL in ALH^RXFP3^ cases (Figure 8E). Notably, the caudal LHb exhibited a patchwork organisation of labelling consistent with the proposed subnuclear structure of the LHb (Quina et al., 2015; F. Wagner et al., 2014). Specifically, fibres were absent in the parvocellular subnucleus of the medial division of the LHb (LHbMPc) and the oval subnucleus of the lateral division of the LHb (LHbLO; Figure 8F).

To determine the precise origin and phenotype of LHb projecting LH/ZI^RXFP3^ cells, we unilaterally targeted a retrograde tracer (pENN.AAV.hSyn.HI.eGFP-Cre.WPRE-SV40; Figure 9A, B) to the LHb and examined the colocalisation of *Rxfp3* with *Slc17a6* (vGlut2) and *Gad1* mRNA transcripts with backlabelled cells in the LH/ZI using RNAscope. 84.9% (± 3.6%) of backlabelled *Rxfp3*+ cells co-expressed *Slc17a6* (Figure 9C, D), while only 13.5% (± 6.5%) co-expressed *Gad1* (Figure 9E). Of the backlabelled *Rxfp3*+ cells co-expressing *Slc17a6*, 91.7% (± 4.0) were located in the IHy or LH (Figure 9F), suggesting the observed glutamatergic ZIR^RXFP3^ input to the LHb likely originates from transfected cells in the IHy or dmLH, rather than the ZIR proper.

**Figure 9.**
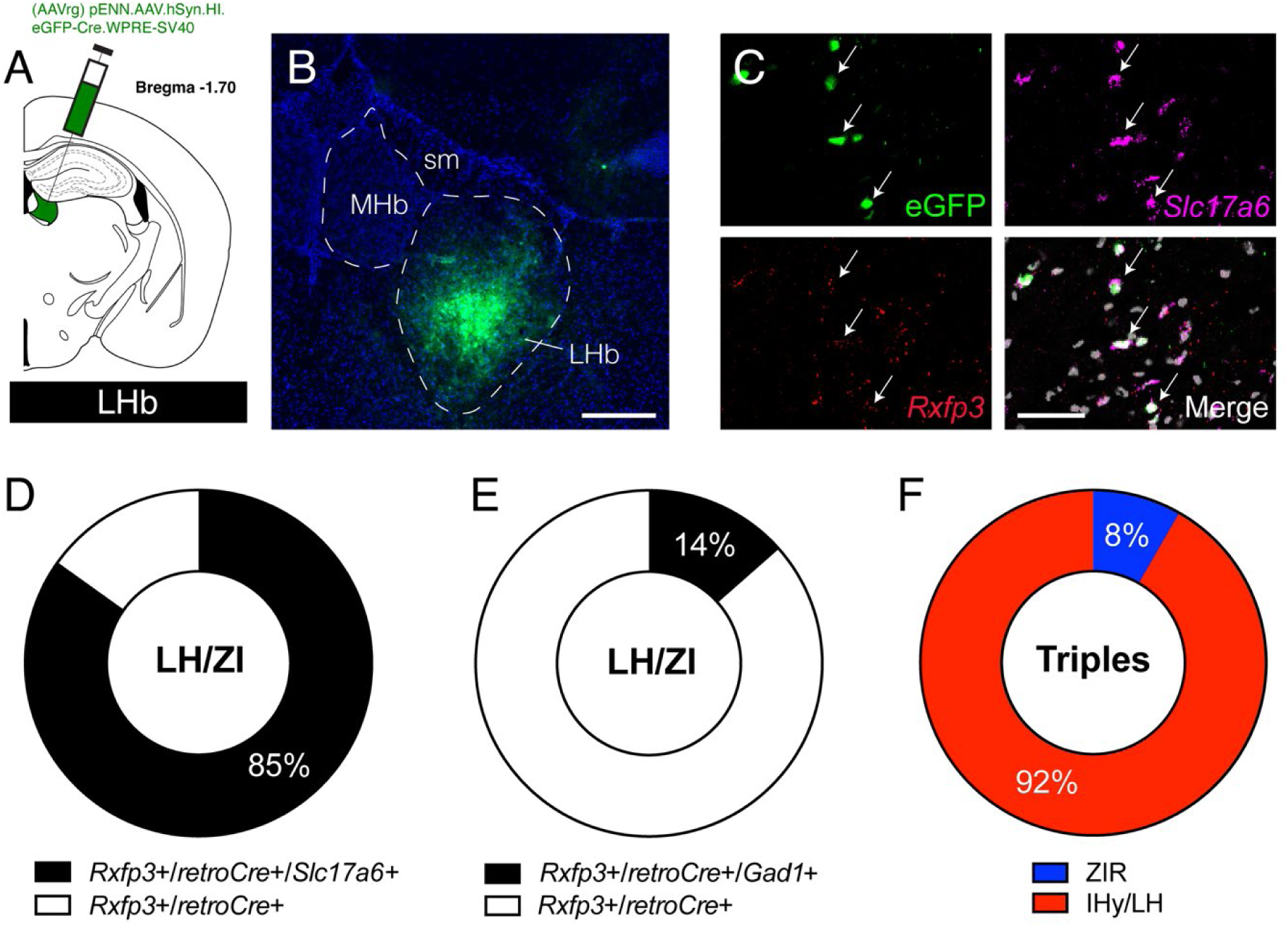
LHb projecting LH/ZI^RXFP3^ cells originate from the LH and are mostly glutamatergic. (A) Retrograde tracing strategy. A retrograde tracer virus was injected into the LHb to trace backlabelled cells in the LH/ZI. (B) Representative stitched fluorescent confocal photomicrograph of retrograde tracer injection site in the LHb. (C) Representative fluorescent confocal photomicrograph of backlabelled cells from an LHb injection in the LH (top left, eGFP) showing co-expression with *Slc17a6* (top right) and *Rxfp3* (bottom left). Merge image shown on the bottom right. (D) Donut graph showing the mean proportion of backlabelled cells from the LHb co-expressing *Rxfp3* only (white) or both *Rxfp3* and *Slc17a6* (black) in the LH/ZI. (E) Donut graph showing the mean proportion of backlabelled cells from the LHb co-expressing *Rxfp3* only (white) or both *Rxfp3* and *Gad1* (black) in the LH/ZI. (F) Donut graph showing the proportion of *Rxfp3*+/*retroCre*+/*Slc17a6*+ cells located in the ZIR (blue) or the IHy/LH (red). MHb, medial habenula; LHb, lateral habenula; sm, stria medullaris. Scale bar in B = 100 µm, Scale bar in C = 50 µm.

##### 3.3.2.6. Dorsal thalamus

The dorsal thalamus was a key target in ZII^RXFP3^ and ZIC^RXFP3^ cases, while ALH^RXFP3^ and ZIR^RXFP3^ cases mostly avoided the area. Rostrally, the Re displayed a moderate density of mGFP+ immunoreactive fibres in both ZII^RXFP3^ and ZIC^RXFP3^ cases (Figure 10A). However, the Re contained a moderate/strong density of mRuby+ boutons in ZIC^RXFP3^ cases, but only contained a low/moderate density of mRuby+ boutons in ZII^RXFP3^ cases, suggesting that most Re input originates from the ZIC rather than the ZII. In ZIC^RXFP3^ cases, moderate to moderate/strong density labelling was observed in intermediate areas of the dorsal thalamus lateral to the LHb, notably in the centrolateral thalamic nucleus (CL), the medial aspect of the laterodorsal thalamic nucleus (LD), and the lateral posterior thalamic nucleus (LP; Figure 10B). Caudally, a dense band of fibres was frequently observed traversing the dorsal aspect of the ventromedial thalamic nucleus (VM), coinciding with the rostral pole of the superior cerebellar peduncle (scp; Figure 10C). Furthermore, in both ZII^RXFP3^ and ZIC^RXFP3^ cases, a moderate to moderate/strong density of dorsoventrally aligned fibres was observed in the ventral portion of the subparafascicular thalamic nucleus (SPF), continuous with labelling in the adjacent PH.

**Figure 10.**
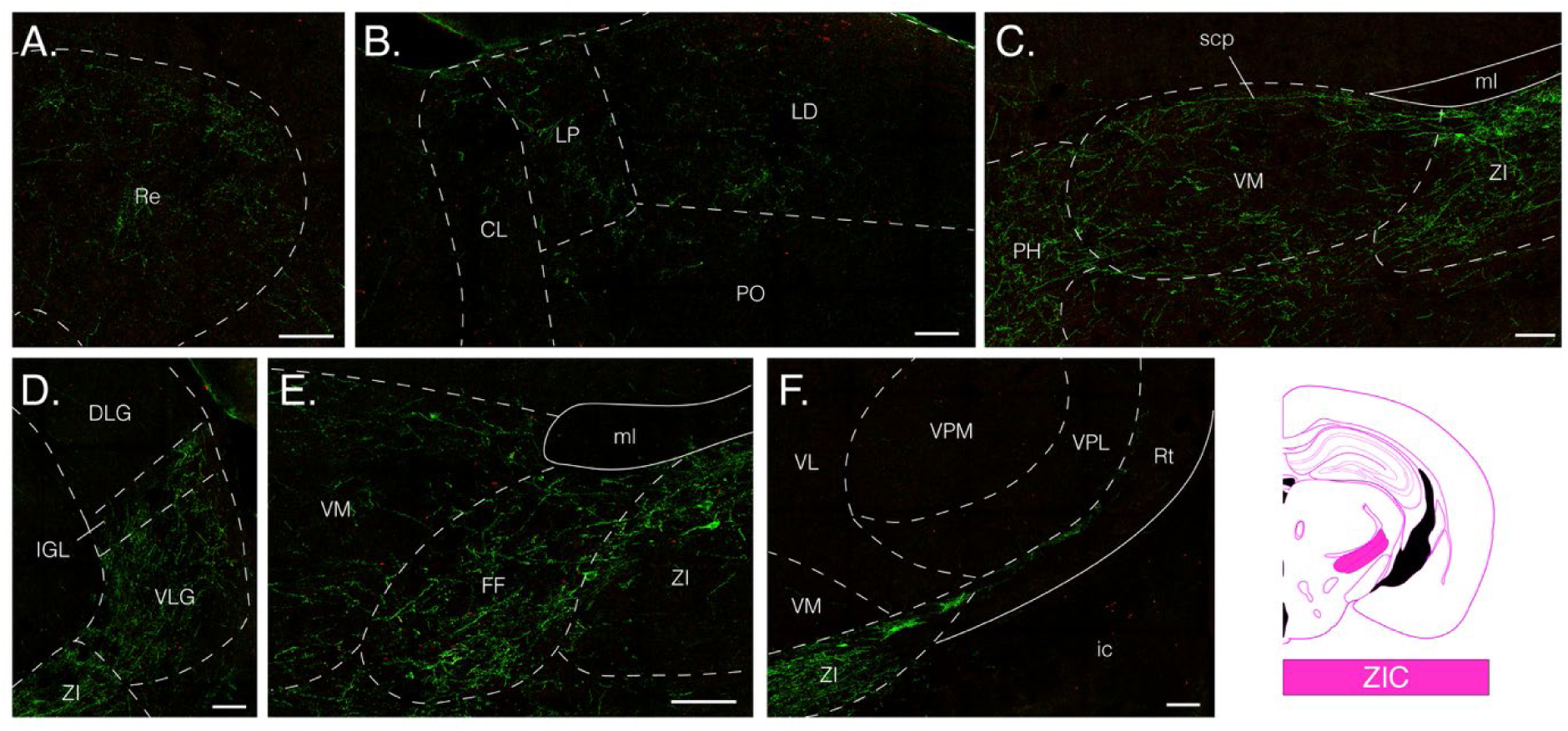
ZIC^RXFP3^ projections to the thalamus. Representative stitched fluorescent confocal photomicrographs of mGFP+ fibres (green) and mRuby+ boutons (red) in the reuniens thalamic nucleus (A), dorsal thalamic nuclei (B), ventromedial thalamic nucleus (C), geniculate nuclei (D), fields of Forel (E), and ventral posterolateral thalamic nucleus/reticular thalamic nucleus border (F). CL, centrolateral thalamic nucleus; FF, fields of Forel; ic, internal capsule; IGL, intergeniculate leaf; LD, laterodorsal thalamic nucleus; LP, lateral posterior thalamic nucleus; ml, medial lemniscus; Po, posterior thalamic nuclear group; PH, posterior hypothalamic area; Re, reuniens thalamic nucleus; Rt, reticular thalamic nucleus; VLG, ventral lateral geniculate nucleus; VL, ventrolateral nucleus of the thalamus; VM, ventromedial thalamic nucleus; VPM, ventral posteromedial thalamic nucleus; VPL, ventral posterolateral thalamic nucleus; ZI, zona incerta;. Scale bars = 100 µm.

##### 3.3.2.7. Zona incerta (ZI)

ALH^RXFP3^ cases did not strongly innervate any subdivision of the ZI (Figure 11A-C), and ZIR^RXFP3^ cases did not strongly innervate intermediate and caudal areas of the ZI (Figure 11D-F). In contrast, ZII^RXFP3^ and ZIC^RXFP3^ cases showed strong innervation of the ZIR and ZIC (Figure 11G-L). These results suggest that ZII^RXFP3^ and ZIC^RXFP3^ cells exhibit inter-sector connectivity, whereas ZIR^RXFP3^ cells do not.

**Figure 11.**
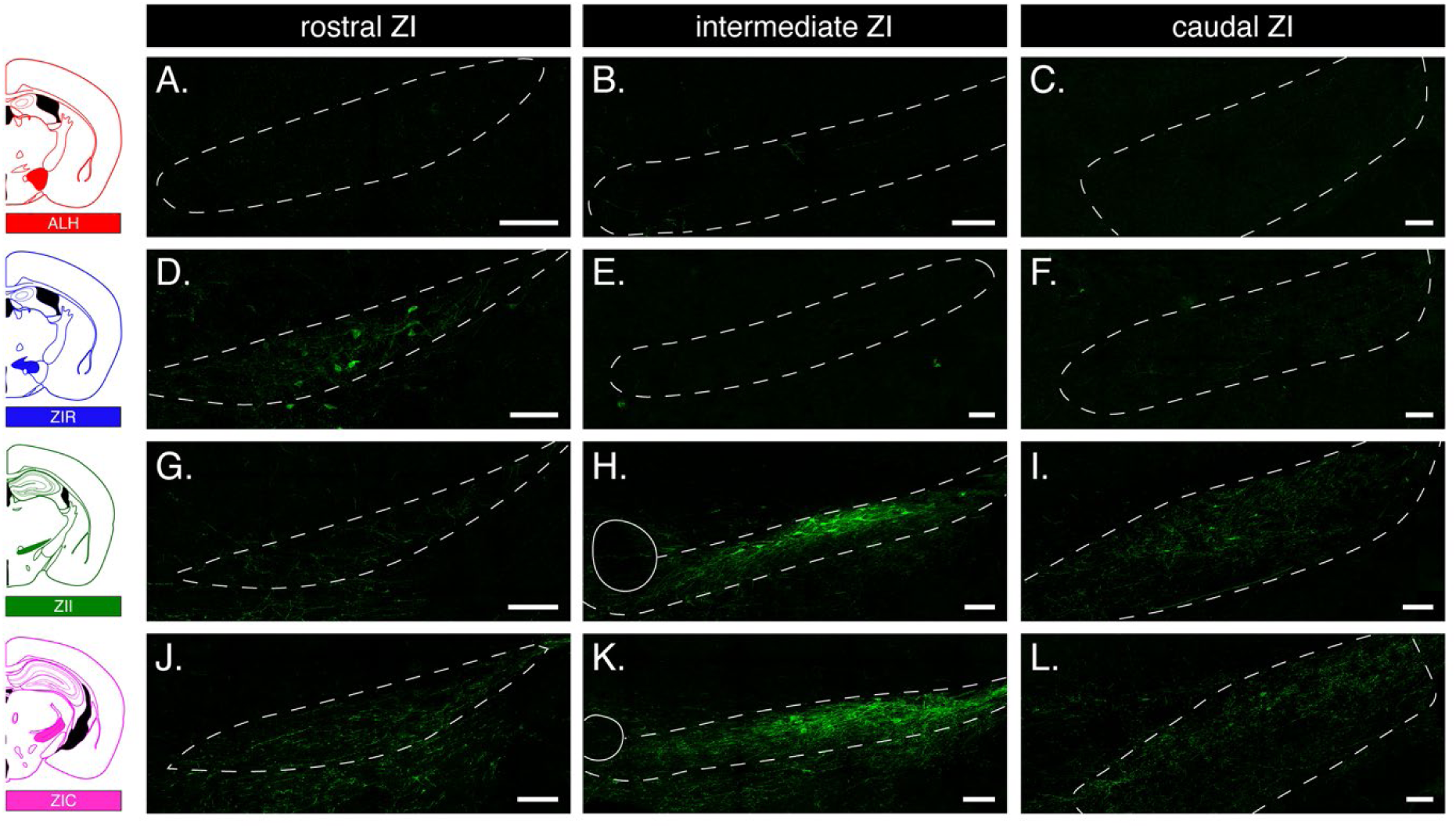
Connectivity within the zona incerta. Representative stitched confocal photomicrographs of mGFP+ immunoreactivity in the rostral ZI (A, D, G, J), intermediate ZI (B, E, H, K), and caudal ZI (C, F, I, L) for an ALH^RXFP3^ case (A-C), a ZIR^RXFP3^ case (D-F), a ZII^RXFP3^ case (G-I), and a ZIC^RXFP3^ case (J-L), demonstrating the interconnectivity within the zona incerta, particularly in ZII^RXFP3^ and ZIC^RXFP3^ cases. Scale bars = 100 µm.

##### 3.3.2.8. Ventral thalamus

Ventral thalamic nuclei were primarily targeted by ZII^RXFP3^ and ZIC^RXFP3^ cases. However, ZIC^RXFP3^ cases generally displayed stronger projections to subregions of the geniculate complex than ZII^RXFP3^ cases. Notably, in ZIC^RXFP3^ cases, strong expression continuous with the lateral ZID/ZIV was observed in the adjacent ventral lateral geniculate nucleus (VLG), and to a lesser extent in the intergeniculate leaf (IGL; Figure 10D). Interestingly, a clear border was observed between the IGL and the dorsal lateral geniculate nucleus, which was devoid of labelling (Figure 10D). In both ZII^RXFP3^ and ZIC^RXFP3^ cases, strong expression continuous with the medial ZID/ZIV was observed in the adjacent fields of Forel (FF; Figure 10E), though mRuby+ boutons were only weakly present in three ZII^RXFP3^ cases. Primarily in ZIC^RXFP3^ cases, a thin band of fibres continuous with expression in the ZIR and rostral ZID/ZIV travelled dorsoventrally and skirted the border of the reticular thalamic nucleus (Rt) and adjacent ventral posterolateral thalamic nucleus (VPL; Figure 10F). For simplicity, expression patterns matching this profile were assigned to the Rt, which accounts for the bulk of Rt expression reported in ZIC^RXFP3^ cases; expression in the Rt proper was generally sparse and comparable to that in other groups.

#### 3.3.3. Mesencephalon

##### 3.3.3.1. Periaqueductal gray (PAG)

The periaqueductal gray was a key target in ZII^RXFP3^ and ZIC^RXFP3^ cases. Rostrally, vertically aligned fibres strongly populated the rostral (RPAG; Figure 12A, B) and supraoculomotor (Su3) divisions. Caudally, both groups displayed similar labelling patterns: the ventrolateral column (VLPAG) received the strongest input, followed by the lateral (LPAG), dorsolateral (DLPAG), and dorsomedial (DMPAG; Figure 12C, D). However, input from ZIC^RXFP3^ cases was consistently stronger than ZII^RXFP3^ input across all PAG columns. Expression was generally stronger in the lateral parts of each column (especially in the LPAG and VLPAG) and decreased in strength closer to the aqueduct (Figure 12C, D).

**Figure 12.**
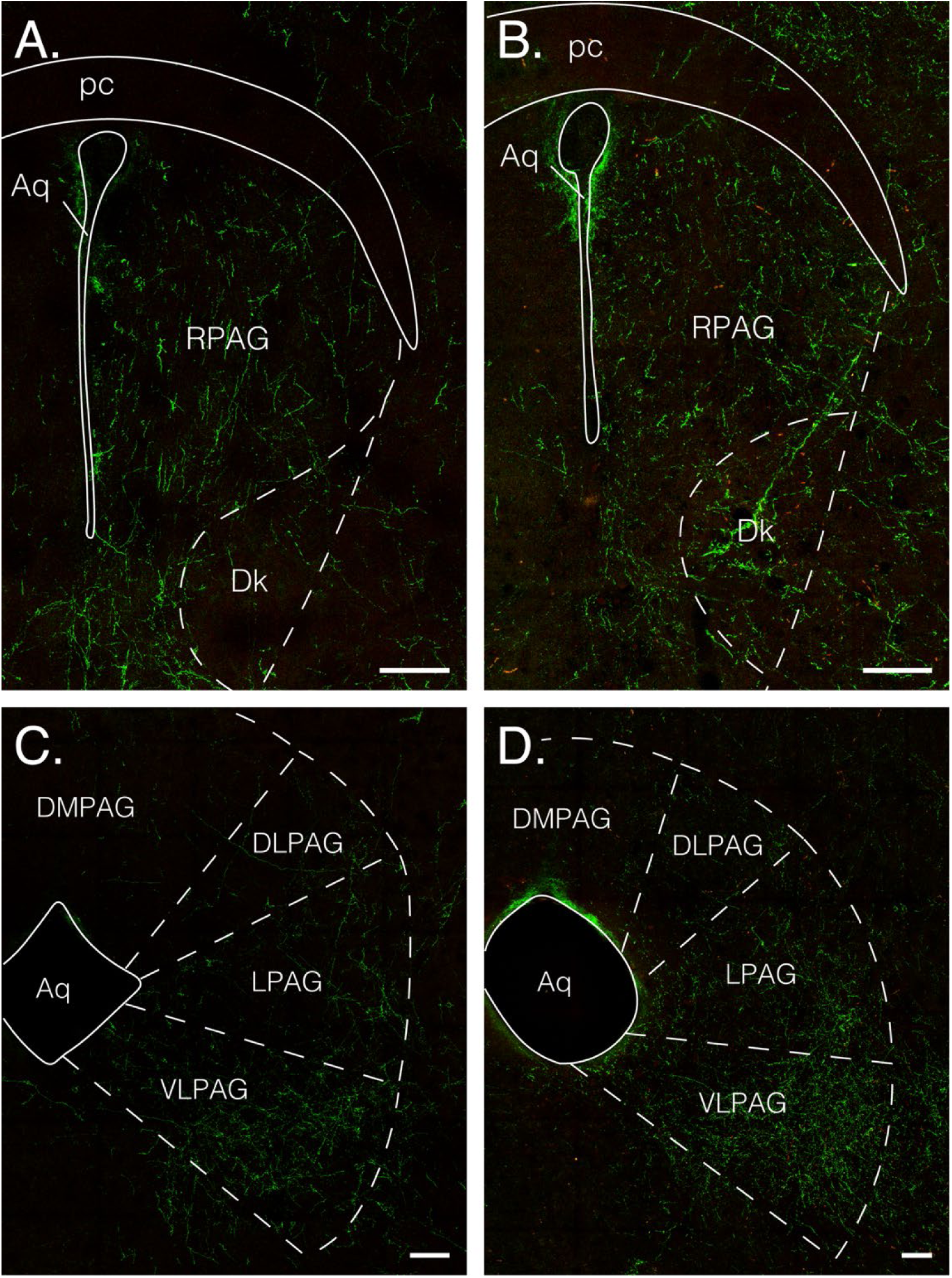
ZII^RXFP3^ cases and ZIC^RXFP3^ cases strongly project to the periaqueductal gray. Representative stitched confocal photomicrographs of mGFP/mRuby expression in the rostral PAG (top) and caudal PAG (bottom) in a ZII^RXFP3^ case (A, C) and a ZIC^RXFP3^ case (B, D). Aq, cerebral aqueduct; Dk, nucleus of Darkschewitsch; DLPAG, dorsolateral periaqueductal gray; DMPAG, dorsomedial periaqueductal gray; LPAG, lateral periaqueductal gray; pc, posterior commissure; RPAG, rostral periaqueductal gray; VLPAG, ventrolateral periaqueductal gray. Scale bars = 100 µm.

Given that the VLPAG was a strong efferent target, we sought to determine the precise origin and phenotype of VLPAG-projecting ZI^RXFP3^ cells. We unilaterally injected a retrograde tracer to the VLPAG (Figure 13A, B) and examined the colocalisation of *Rxfp3* with *Slc17a6* (vGlut2) and *Gad1* mRNA transcripts with backlabelled cells in the ZI. 89.1% (± 2.1%) of backlabelled ZI *Rxfp3*+ cells co-expressed *Gad1* (Figure 13C, E), while only 8.1% (± 2.3 %) co-expressed *Slc17a6* (Figure 13D). Most backlabelled *Rxfp3*+/*Gad1*+ cells were observed in the ZIV (90.2% ± 4.6%) and not the ZID (Figure 13F).

**Figure 13.**
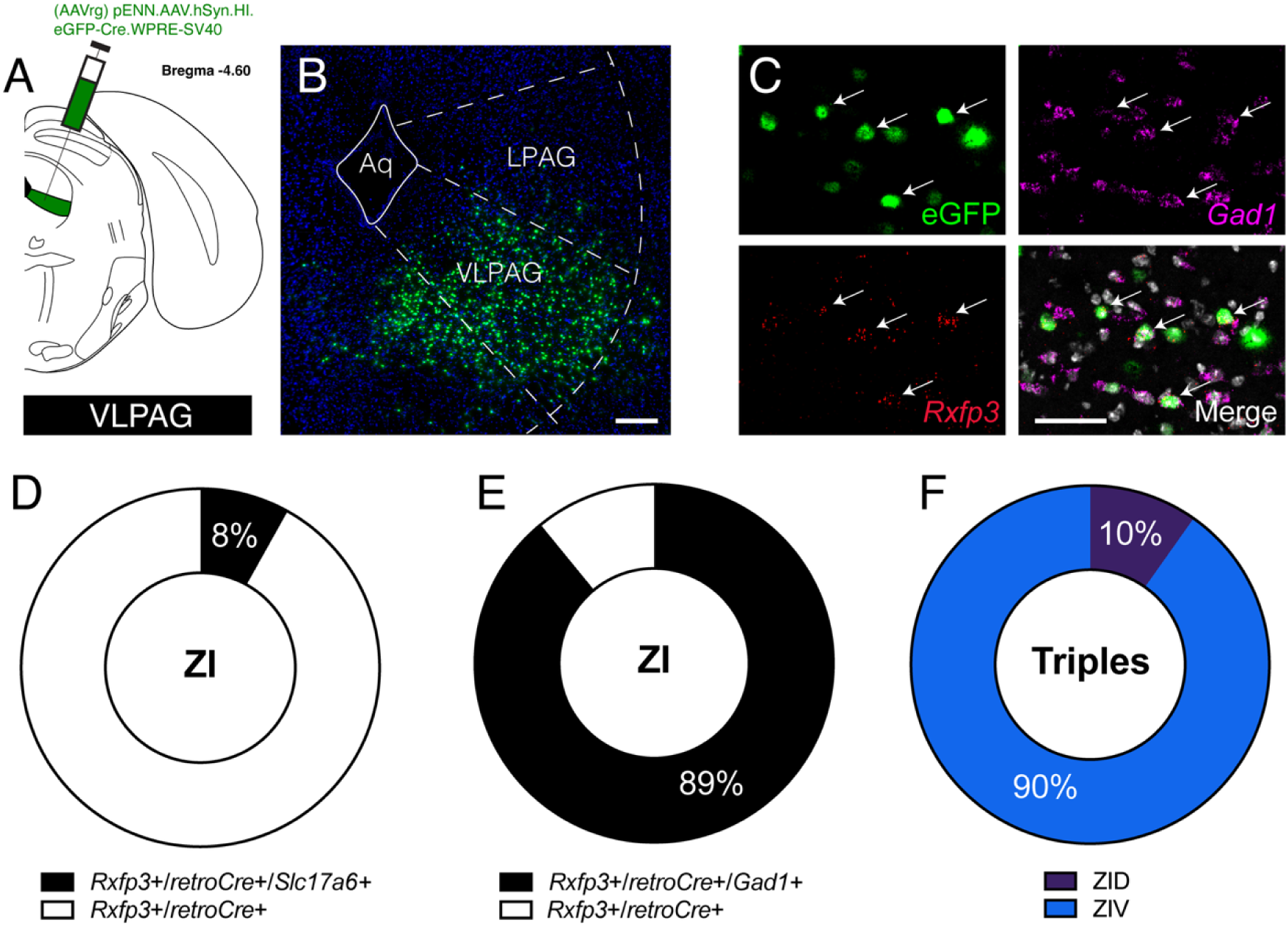
VLPAG projecting ZI^RXFP3^ cells are mostly GABAergic. (A) Retrograde tracing strategy. A retrograde tracer virus was injected into the VLPAG to trace backlabelled cells in the ZI. (B) Representative stitched fluorescent confocal photomicrograph of retrograde tracer injection site in the VLPAG. (C) Representative fluorescent confocal photomicrograph of backlabelled cells from a VLPAG injection in the ZI (top left, eGFP) showing co-expression with *GAD1* (top right) and *Rxfp3* (bottom left). Merge image shown on the bottom right. (D) Donut graph showing the mean proportion of backlabelled cells from the VLPAG co-expressing *Rxfp3* only (white) or both *Rxfp3* and *Slc17a6* (black) in the ZI. (E) Donut graph showing the mean proportion of backlabelled cells from the VLPAG co-expressing *Rxfp3* only (white) or both *Rxfp3* and *Gad1* (black) in the ZI. (F) Donut graph showing the proportion of *Rxfp3*+/*retroCre*+/*Gad1*+ cells located in the ZID (purple) or the ZIV (blue). Aq, cerebral aqueduct; LPAG, lateral periaqueductal gray; VLPAG, ventrolateral periaqueductal gray. Scale bar in B = 100 µm, Scale bar in C = 50 µm.

##### 3.3.3.2. Tegmentum

Many tegmental areas were key targets of ZII^RXFP3^ and ZIC^RXFP3^ cases, with ZIC^RXFP3^ cases consistently producing stronger innervation patterns than ZII^RXFP3^ cases across most analysed tegmental regions. Rostrally, fibres continuous with the dorsal part of the RPAG populated the nucleus of the posterior commissure (PCom; Figure 14A), precommissural nucleus (PrC), and retroparafascicular nucleus (RPF). Fibres mainly occupied the medial part of the anterior pretectal nucleus (APT) and indiscriminately occupied the medial (MPT), posterior (PPT), and olivary (OPT) pretectal nuclei in ZIC^RXFP3^ cases (Figure 14A), whereas ZII^RXFP3^ cases weakly targeted these areas. Regions immediately ventral to the RPAG also received moderate/strong input, including the nucleus of Darkschewitsch (Dk), interstitial nucleus of Cajal (InC), and Edinger-Westphal nucleus (EW). In intermediate areas of the tegmentum, diagonally oriented fibres sparsely occupied the dorsal part of the midbrain reticular nucleus (MRN) but densely clustered around the central part of the nucleus. A thick band of fibres traversed the lateral border of the red nucleus (R) and invaded the medial MRN (Figure 14B). Additionally, a separate band of moderate-density fibres travelled through the dorsal MRN and terminated in the sagulum nucleus (Sag). In ventral parts of the tegmentum, a dense fibre cluster was observed in a small ventromedial part of the substantia nigra, compact part (SNC; Figure 14B), a nucleus otherwise devoid of expression. Furthermore, low-density labelling was observed throughout the ventral tegmental area (VTA). Caudally, strong labelling continuous with the lateral part of the VLPAG occupied the medial aspect of the cuneiform nucleus (CnF) across its rostrocaudal extent (Figure 14D). Both the ventral tegmental nucleus (VTg) and dorsal raphe nucleus (DR) received moderate input.

**Figure 14.**
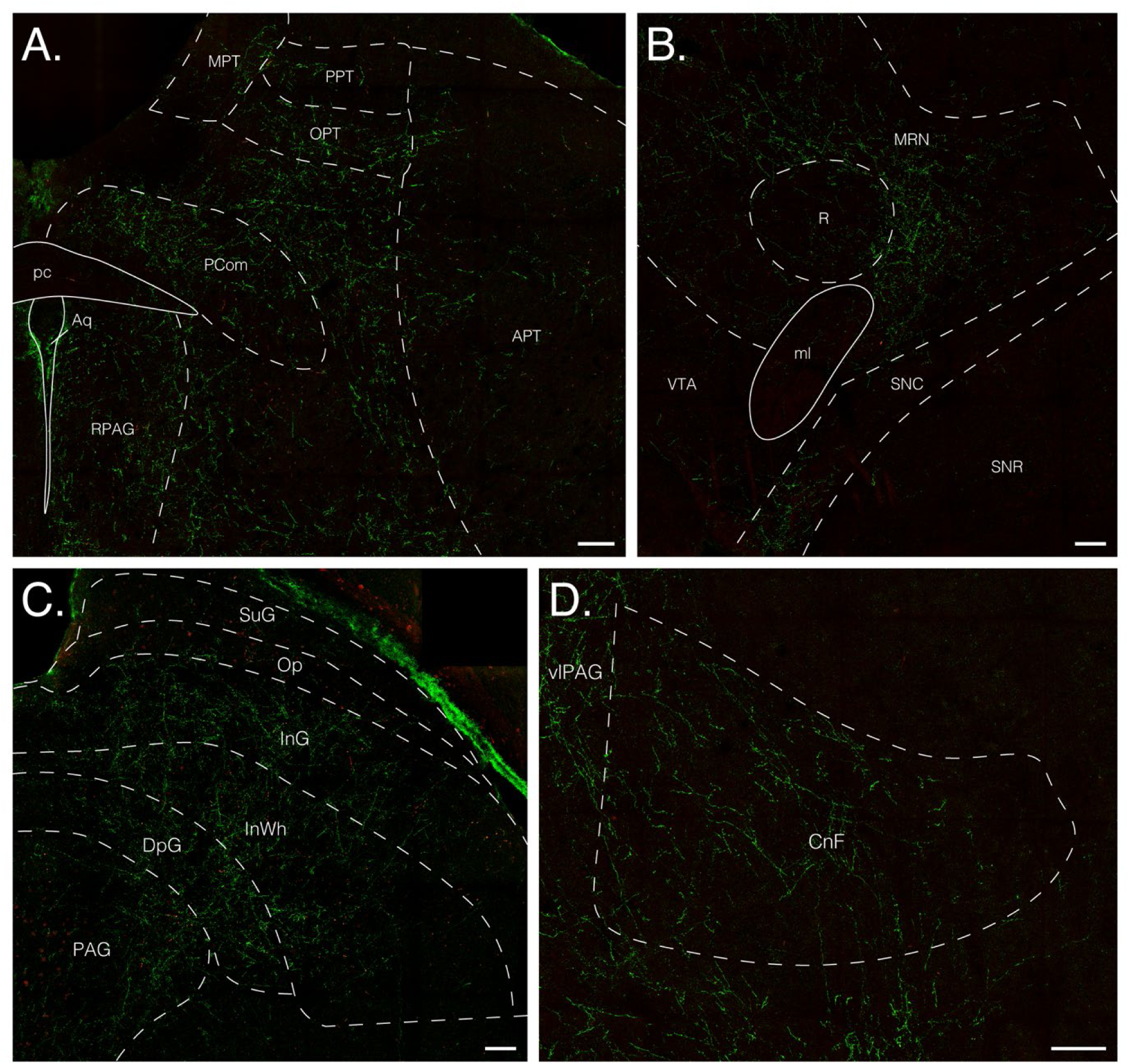
ZIC^RXFP3^ projections to the mesencephalon. Representative stitched confocal photomicrographs of mGFP/mRuby expression in various subnuclei of the pretectal area, rostral periaqueductal gray, and nucleus of the posterior commissure (A), midbrain reticular nucleus and red nucleus (B), superior colliculus (C), and cuneiform nucleus (D). APT, anterior pretectal nucleus; Aq, cerebral aqueduct; CnF, cuneiform nucleus; DpG, deep gray layer of the superior colliculus; InG, intermediate gray layer of the superior colliculus; InWh, intermediate white layer of the superior colliculus; ml, medial lemniscus; MPT, medial pretectal nucleus; MRN, midbrain reticular nucleus; Op, optic nerve layer of the superior colliculus; PAG, periaqueductal gray; pc, posterior commissure; PCom, nucleus of the posterior commissure; PPT, posterior pretectal nucleus; R, red nucleus; RPAG, rostral periaqueductal gray; SNC, substantia nigra, compact part; SNR, substantia nigra, reticular part; SuG, superficial gray layer of the superior colliculus; VTA, ventral tegmental area. Scale bars = 100 µm.

##### 3.3.3.3. Superior colliculus

ZIC^RXFP3^ cases produced the strongest labelling in subregions of the superior colliculus (Figure 14C). Generally, expression was moderate in the medial parts of the ventral subnuclei: the deep gray layer (DpG), intermediate gray layer (InG), and intermediate white layer (InWh). Occasional fibres were observed in the more lateral parts of these nuclei.

Low/moderate labelling was evident in dorsal nuclei: the optic nerve layer (Op) and superficial gray layer (SuG).

#### 3.3.4. Rhombencephalon

##### 3.3.4.1. Pons

In both ZIC^RXFP3^ and ZII^RXFP3^ cases, the pontine reticular nucleus, oral part (PnO) received massive input along its rostrocaudal extent (Figure 15A). Indeed, mGFP+ labelling in the PnO accounted for 71.5% (± 2.3%) of pontine input and 28.3% (± 3.2%) of overall input in ZII^RXFP3^ cases, and 56.5% (± 3.1%) of pontine input and 21.5% (± 0.9%) overall input in ZIC^RXFP3^ cases (Figure 15B). In the rostral pons, dense fibre bands decussated from the PnO to innervate the median raphe nucleus (MnR), reticulotegmental nucleus (RtTg), and the lateral border of the pedunculopontine tegmental nucleus (PPTg; Figure 15C). In the intermediate pons, PnO expression became continuous with expression in the pontine reticular nucleus, caudal part (PnC), and the nucleus raphe pontis (RPO). Caudally, strong labelling was observed in the ventral half of the nucleus incertus (NI), continuous with strong expression in the central gray of the pons (CGPn; Figure 15D). Interestingly, labelling generally avoided the dorsal tegmental nucleus (DTg) and laterodorsal tegmental nucleus (LDTg) embedded within the CGPn (Figure 15D). Moderate density labelling was observed in the subcoeruleus nucleus (SubC) and sublaterodorsal nucleus (SLD).

**Figure 15.**
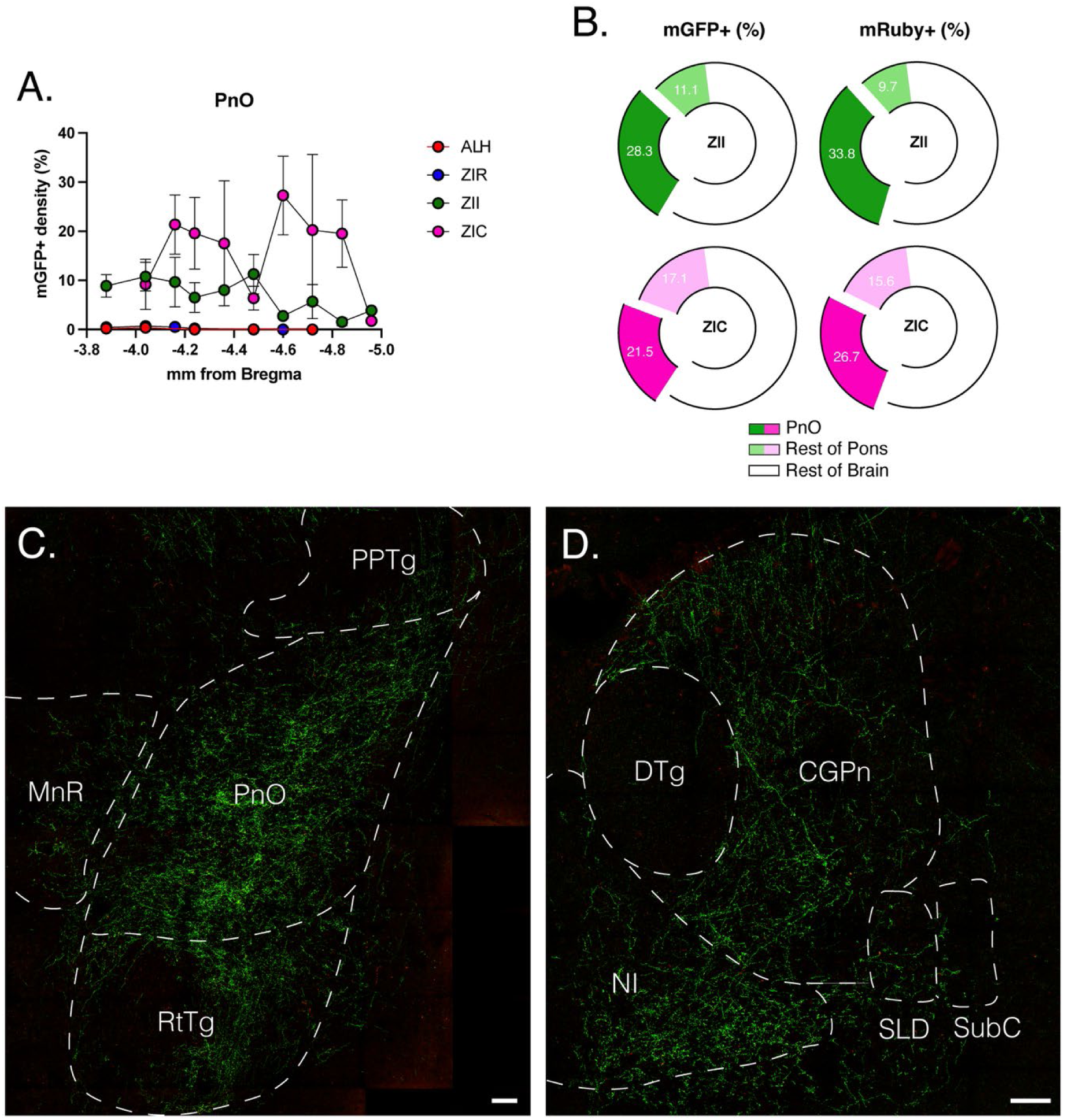
Projections to the pons. (A) Graph showing the average mGFP+ density in the PnO separated by Bregma level for each injection site group. (B) Donut graphs showing the average relative percentage of mGFP+ fibres (left) and mRuby+ boutons (right) observed in the PnO (dark green, dark pink) and the rest of the pons (light green, light pink) in both ZII^RXFP3^ cases (green) and ZIC^RXFP3^ cases (pink) as a proportion of total observed mGFP+/mRuby observed throughout the entire brain. (C, D) Representative stitched confocal photomicrographs of mGFP/mRuby expression in the rostral pons (C) and caudal pons (D). CGPn, central gray of the pons; DTg, dorsal tegmental nucleus; MnR, median raphe nucleus; NI, nucleus incertus; PnO, pontine reticular nucleus, oral part; PPTg, pedunculopontine tegmental nucleus; RtTg, reticulotegmental nucleus; SLD, sublaterodorsal nucleus; SubC, subcoeruleus nucleus. Scale bars = 100 µm.

##### 3.3.4.2. Medulla

Low to low/moderate labelling was observed in the medulla for ZII^RXFP3^ and ZIC^RXFP3^ cases. However, moderate/strong labelling was observed in the magnocellular reticular nucleus (MARN), abducens nucleus (6N), and raphe magnus nucleus (RMg). Moderate labelling was also observed in the rostral part of the gigantocellular reticular nucleus (Gi).

#### 3.3.5. White matter

In cases with strong LHb expression (mainly ZIR^RXFP3^ case 172), fibres were frequently observed in the fasciculus retroflexus (fr), continuous with expression in the caudal part of the LHb. In ZIR^RXFP3^ cases, consistent moderate/strong labelling was observed along the borders of the stria medullaris of the thalamus (sm). In ZIC^RXFP3^ cases, a strong density of dorsoventrally aligned fibres occupied the trapezoid body (tz), infiltrating the dorsal part of the pyramidal tract (py) at the level of the Gi. In ZII^RXFP3^ and ZIC^RXFP3^ cases, moderate to moderate/intense labelling was observed in the medial lemniscus (ml) at the level of the VTA, often continuous with expression in the ventral MRN (Figure 12B).

## 4. Discussion

In this study, we injected a Cre-dependent anterograde tracer virus into four distinct sites of the LH/ZI in RXFP3-Cre mice and analysed their efferent connectivity patterns throughout the entire brain. Injection cases were grouped based on their viral spread properties: ALH^RXFP3^, where spread was contained to the anterior third of the LH; ZIR^RXFP3^, where spread infiltrated the incertohypothalamic area and dorsomedial LH; ZII^RXFP3^, where spread was primarily contained to the rostral half of the ZID/ZIV; and ZIC^RXFP3^, where spread was primarily contained to the caudal half of the ZID/ZIV. At the macroscale level, we demonstrated that ALH^RXFP3^ and ZIR^RXFP3^ cells primarily project to regions within the diencephalon, while ZII^RXFP3^ and ZIC^RXFP3^ cells predominantly project downstream to midbrain and pontine nuclei. At the individual nucleus level, we demonstrated that each injection site group produced unique projection patterns, particularly to nuclei implicated in threat and defensive behaviour. This supports a previous finding from our lab where chemogenetically activating a large population of LH/ZI^RXFP3^ cells during conditioned fear retrieval produced multiple behavioural phenotypes, including increased locomotion and escape-like jumping behaviour (Richards et al., 2025). However, the previous study examined only male mice, whereas the current study assessed both sexes. Therefore, future work should functionally interrogate female mice to determine if they display analogous phenotypes. Nevertheless, our results suggest that LH/ZI^RXFP3^ cells exhibit distinct efferent projection patterns throughout the brain depending on their topographical location within these nuclei, likely reflecting the functional diversity of these neurons.

### 4.1. Projections to the dorsal premammillary nucleus

The PMD was a key target in ALH^RXFP3^ cases, ZIR^RXFP3^ cases, and one ZII^RXFP3^ case with transfected cells in the LH. Multiple lines of evidence suggest that PMD efferents originate from the LH. On the other hand, we found that the major source of PMD projections was the ZIR^RXFP3^ cells. This discrepancy may be due to the experimental approach adopted in our study. For example, in rodents, a projection from the LH to the PMD is well-documented (Comoli et al., 2000; Faturi et al., 2014; Goto et al., 2005; Hahn & Swanson, 2012; Viellard et al., 2024), while only sparse projections to the PMD have been reported from ZI A13 dopamine cells in one recent study (Bono et al., 2025). Furthermore, reported efferents to the PMD originate from both the juxtadorsomedial LH and the suprafornical LH (Faturi et al., 2014; Hahn & Swanson, 2012; Viellard et al., 2024) in the rat. Although undefined in common mouse atlases, these regions appear to overlap with the IHy area defined in our study, where we observed dense populations of mGFP+ immunoreactive cell bodies in ZIR^RXFP3^ cases. However, to definitively identify the exact origin of LH/ZI^RXFP3^ inputs to the PMD, a Cre-dependent retrograde tracer should be injected into the PMD of RXFP3-Cre mice.

The PMD is a small hypothalamic nucleus mainly consisting of cholecystokinin-expressing glutamatergic neurons that are activated by various threats, including carbon dioxide exposure (Johnson et al., 2011), predator exposure (Melleu et al., 2022; Mendes-Gomes et al., 2020), and social defeat stress (De Almeida et al., 2022; Faturi et al., 2014). Of particular interest, activating the PMD induces context-specific escape behaviours (Laing et al., 2023; W. Wang et al., 2021). Given that we have demonstrated that chemogenetic activation of LH/ZI^RXFP3^ cells induces panic-like jumps in a fear conditioning chamber (where escape is impossible) in some mice, it is possible that downstream PMD cells were activated by glutamatergic LH^RXFP3^ cells to permit context-appropriate escape – i.e. jumping to avoid the grid floor. Future studies should examine the neurochemical phenotype of LH^RXFP3^ neurons that project to the PMD and manipulate this pathway across different threatening contexts to determine its role in context-specific escape behaviour.

### 4.2. Projections to the lateral habenula

Previously, we demonstrated that LH/ZI^RXFP3^ cells projected to the LHb, but did not determine the precise origin of these projections or detail their intra-LHb innervation patterns (Richards et al., 2025). In the current study, we discovered that ALH^RXFP3^ and ZIR^RXFP3^ neurons strongly project to the LHb but innervate distinct LHb territories. ALH^RXFP3^ cases mainly project to the LHbL and selectively avoid the LHbLO and LHbMPc subnuclei, while ZIR^RXFP3^ cases primarily project to the LHbM. Additionally, retrograde tracing and neurochemical phenotyping revealed that most LHb input originated from a subset of *Rxfp3*+/*Slc17a6*+ cells throughout the LH, and not from *Rxfp3*+ ZIR cells. Overall, our results indicate that LHb projecting LH^RXFP3^ cells are a subset of topographically organised glutamatergic neurons, where those in the dmLH preferentially project to the LHbM, while those in the ALH proper preferentially project to the LHbL.

Our findings largely echo a recent study demonstrating that the LH contains multiple, topographically distinct, glutamatergic subtypes with unique projection patterns to the LHb (Calvigioni et al., 2023). Of relevance to the current study, they demonstrated that LHb-projecting *Esr1*+ LH neurons populate the dorsomedial LH, innervate the LHbM, and specifically avoid projecting to the LHbMPc and LHbLO, precisely mirroring the properties of dmLH^RXFP3^ neurons. Additionally, they demonstrated that LHb-projecting neuropeptide-Y+ LH neurons populate the ALH proper and predominantly terminate in the LHbL, mirroring the properties of ALH^RXFP3^ neurons. It therefore seems likely that dmLH^RXFP3^ neurons are a subset of *Esr1*+ LH neurons, and ALH^RXFP3^ cells are a subset of neuropeptide-Y+ LH neurons, however future studies are needed to verify this hypothesis.

Converging studies have shown that glutamatergic LH-LHb neurons encode aversion, however these neurons are typically studied as a homogeneous population, (Lazaridis et al., 2019; Lecca et al., 2017; Zheng et al., 2022). On the other hand, recent electrophysiological studies have shown that in response to footshock, LHbL neurons show excitation, while LHbM neurons show inhibition (Congiu et al., 2019, 2023). Therefore, our discovery that topographically distinct LH^RXFP3^ cells selectively innervate either the LHbL or LHbM implies that these subpopulations may make distinct contributions to aversive processing. Furthermore, the aforementioned *Esr1*+ and neuropeptide-Y+ LHb-projecting glutamatergic neurons each play distinct roles in aversion, with the former driving real-time place aversion, and the latter driving unsupported rearing behaviour (Calvigioni et al., 2023), suggesting that such a functional opposition between discrete LH^RXFP3^ subsets would not be unprecedented.

### 4.3. Projections to the periaqueductal gray

We demonstrated that the LPAG/VLPAG were strongly innervated by ZII^RXFP3^ and ZIC^RXFP3^ cells, but not by ALH^RXFP3^ and ZIR^RXFP3^ cells. Furthermore, retrograde tracing and neurochemical phenotyping revealed that most VLPAG input arose from a subset of *Gad1*+ neurons in the ZID/ZIV. Taken together, we have shown that a topographically defined subset of GABAergic ZI^RXFP3^ neurons projects to the LPAG/VLPAG.

GABAergic ZI neurons project extensively throughout the PAG (Ahmadlou et al., 2021; Liu et al., 2017; Tong et al., 2025; Venkataraman et al., 2019; Yu et al., 2021; Zhao et al., 2019) and again are frequently interrogated as a homogeneous population. However, GABAergic ZI neurons consist of neurochemically defined subsets that express tyrosine hydroxylase (Negishi et al., 2020; Venkataraman et al., 2021), tachykinin-1 (Ahmadlou et al., 2021), somatostatin (Z. Li et al., 2021), parvalbumin (Wallén-Mackenzie et al., 2020) and many others (V. Cheung et al., 2021; Z. Li et al., 2021; Liu et al., 2017; Zhu et al., 2025), some of which target discrete PAG columns. For example, tachykinin-1+ ZI neurons target the LPAG/VLPAG (Ahmadlou et al., 2021), calretinin+ ZI neurons target the DMPAG (Z. Li et al., 2021), and parvalbumin+ ZI neurons terminate along the lateral border of the DLPAG/LPAG/VLPAG (H. Wang et al., 2020; Zhou et al., 2018). Here we showed that RXFP3+ GABAergic ZI neurons specifically innervate the LPAG/VLPAG, similar to tachykinin-1+ ZI neurons. Moreover, GABAergic ZI neurons display topographically arranged projections to the PAG: the medial ZI innervates the LPAG/VLPAG, the lateral ZI innervates the DLPAG, and the ZIR innervates the DMPAG (Yang et al., 2022). Combined with the canonical view that different PAG columns have distinct functional roles (Reis et al., 2023; Zhang et al., 2024), it is clear that the GABAergic ZI-PAG pathway should not be treated as a single, homogenous entity. Future research should focus on examining neurochemically and topographically distinct GABAergic ZI-PAG populations to parse their distinct roles.

The strong projections observed in the LPAG/VLPAG from ZII^RXFP3^ and ZIC^RXFP3^ cells suggest that this pathway may regulate the expression of defensive behaviours. It is widely understood that activation of the LPAG/VLPAG causes defensive freezing behaviour (Fanselow et al., 1995; La-Vu et al., 2022), while activating the dorsal PAG evokes panic-like jumping and escape (Deng et al., 2016; Evans et al., 2018). Recent evidence suggests that VLPAG-mediated freezing arises from activity of vGlut2+ expressing neurons in the area; however, GAD2+ VLPAG neurons can locally suppress vGlut2+ VLPAG neurons to inhibit freezing (Tovote et al., 2016), and activating a sparse population of cholecystokinin+/vGlut2+ VLPAG neurons can drive escape behaviours (La-Vu et al., 2022). Therefore, GABAergic ZII/ZIC^RXFP3^ cells may regulate either passive freezing or active escape behaviours in response to threats, depending on their specific, neurochemically defined VLPAG targets, which remain to be determined. This pathway may also regulate fear learning rather than just fear expression, as chemogenetic inhibition of the VLPAG impairs the acquisition of conditioned suppression of reward (Arico et al., 2017). As this behaviour is not dependent on PAG activity for its expression (Amorapanth et al., 1999), this suggests that the impediment is due to impaired associative learning rather than fear expression.

### 4.4. Other key projections

At a macroscale level, ALH^RXFP3^ and ZIR^RXFP3^ cases generally innervated areas within the diencephalon and exhibited unique upstream projections. In particular, ZIR^RXFP3^ cases uniquely innervated areas of the basal forebrain (LS, MS, DB, SI) and the preoptic hypothalamus (LPO, MPO). Interestingly, studies that have demonstrated a functional role of LH to basal forebrain or preoptic nuclei have interrogated melanin-concentrating hormone-or orexin-expressing LH neurons (De Luca et al., 2022; Jego et al., 2013; C. Ma et al., 2023), which do not express RXFP3 (Richards et al., 2025). Although one study has demonstrated that GABAergic LH projections to the diagonal band drive feeding behaviour and reduce anxiety (Cassidy et al., 2019), no other studies have functionally interrogated these pathways. Given that the many neuroanatomical tract-tracing studies, including ours, provide evidence of connectivity, future studies should attempt to parse the function of these connections.

Conversely, ZII^RXFP3^ and ZIC^RXFP3^ cases primarily exhibited downstream projections to several mesencephalic and rhombencephalic regions heavily implicated in arousal, including the PPTg, NI, and CGPn (Dugan et al., 2023; Kroeger et al., 2022; S. Ma et al., 2017; Ryan et al., 2011; Wei et al., 2024). Of note were the particularly dense projections to the PnO, comprising approximately a quarter of the overall output of ZII^RXFP3^ and ZIC^RXFP3^ cases.

Although a GABAergic ZI-PnO pathway is well-established, only a handful of studies have explored its function in different domains (Ahmadlou et al., 2021; Zhao et al., 2019; Zhu et al., 2025). Optogenetically activating the terminals of Pde11a+ GABAergic ZI neurons in the PnO promotes wakefulness, suggesting that a subset of GABAergic ZI neurons increases arousal levels by inhibiting the PnO (Zhu et al., 2025). Therefore, it is possible that GABAergic ZII/ZIC^RXFP3^ cells may regulate PnO activity to promote increased arousal levels based on environmental demands. Combined with the finding that ZII^RXFP3^ and ZIC^RXFP3^ cases also projected strongly to regions implicated in generating panic-like defensive behaviours, including the SC, CnF, and PH (Biagioni et al., 2012; Bindi et al., 2023; Caggiano et al., 2018; Da Silva Soares et al., 2019; Falconi-Sobrinho et al., 2017), ZII/ZIC^RXFP3^ projections to arousal-promoting regions may be necessary for an organism to generate active defensive responses to immediate threats.

### 4.5. Conclusion

This study is the first to demonstrate hodological variability within a relatively continuous RXFP3+ population in the ZI and LH. Future studies should take this into account when examining the connectivity profile of RXFP3+ neurons in other RXFP3-dense areas of the brain (e.g. BST, LS), rather than assuming hodological uniformity. This hodological variability is likely to translate into functional variability, which warrants further interrogation.

## Acknowledgements

We thank the Central Animal Facility staff at Macquarie University for animal husbandry.

## Contributions

BKR and CJP designed the experiment. BKR performed all experiments and wrote the manuscript. AIJK assisted with mouse brain registration and JLC assisted with data interpretation. JHK, AJL, CJP acquired funds for the research. All authors reviewed and edited the manuscript.

## Data availability statement

The data that support the findings of this study are available from the corresponding author upon reasonable request.

