## Supplementary materials for "Efferent projections of topographically distinct relaxin family peptide receptor-3 (RXFP3) lateral hypothalamus/zona incerta cells"

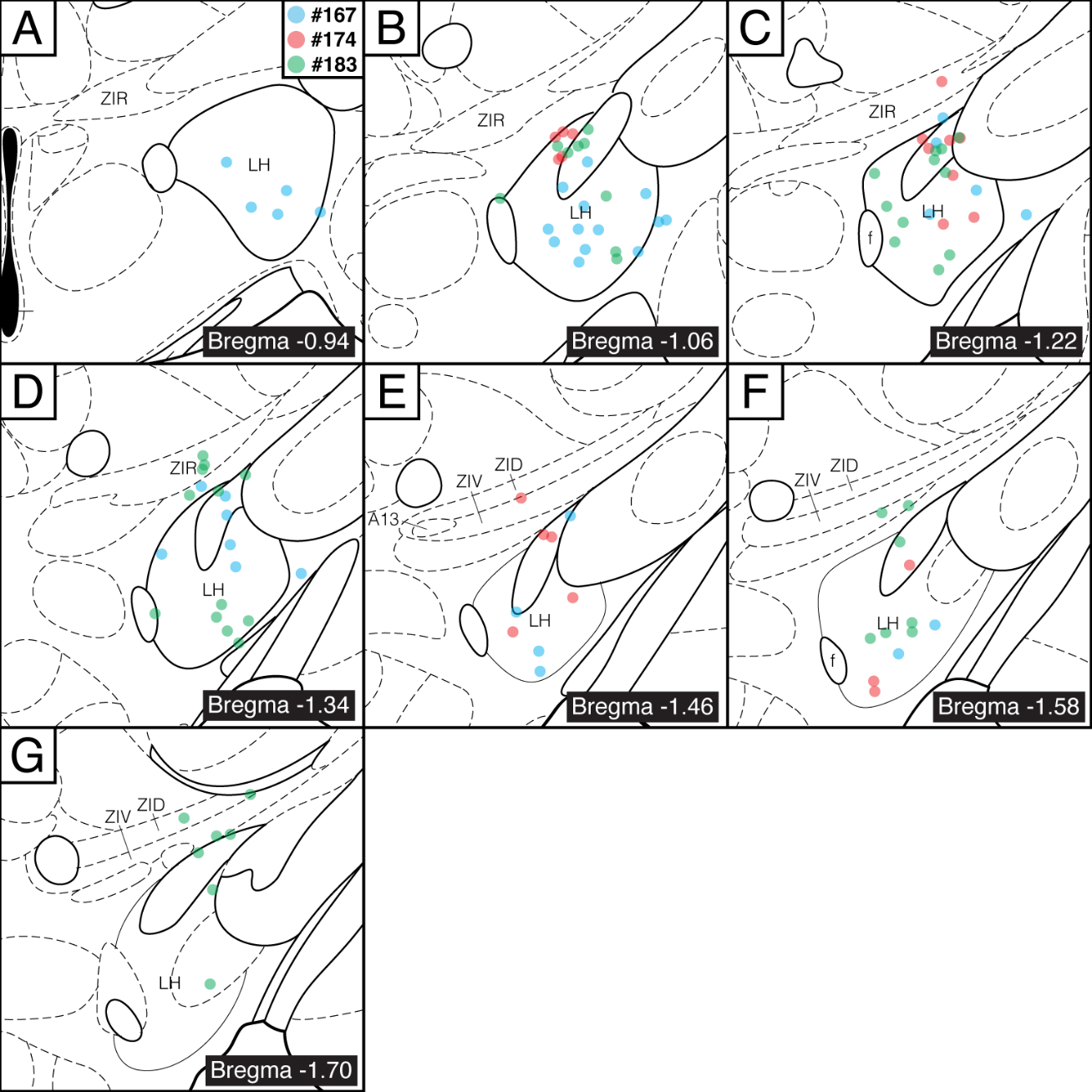

**Supplementary Figure 1.** Coronal brain schematics depicting anterograde tracer injection sites in the ALH^RXFP3^ group of RXFP3-Cre mice (*n* = 3) mapped onto plates from the Mouse Brain in Stereotaxic Coordinates (Paxinos & Franklin, 2004). Each dot represents an mGFP immunoreactive cell body.

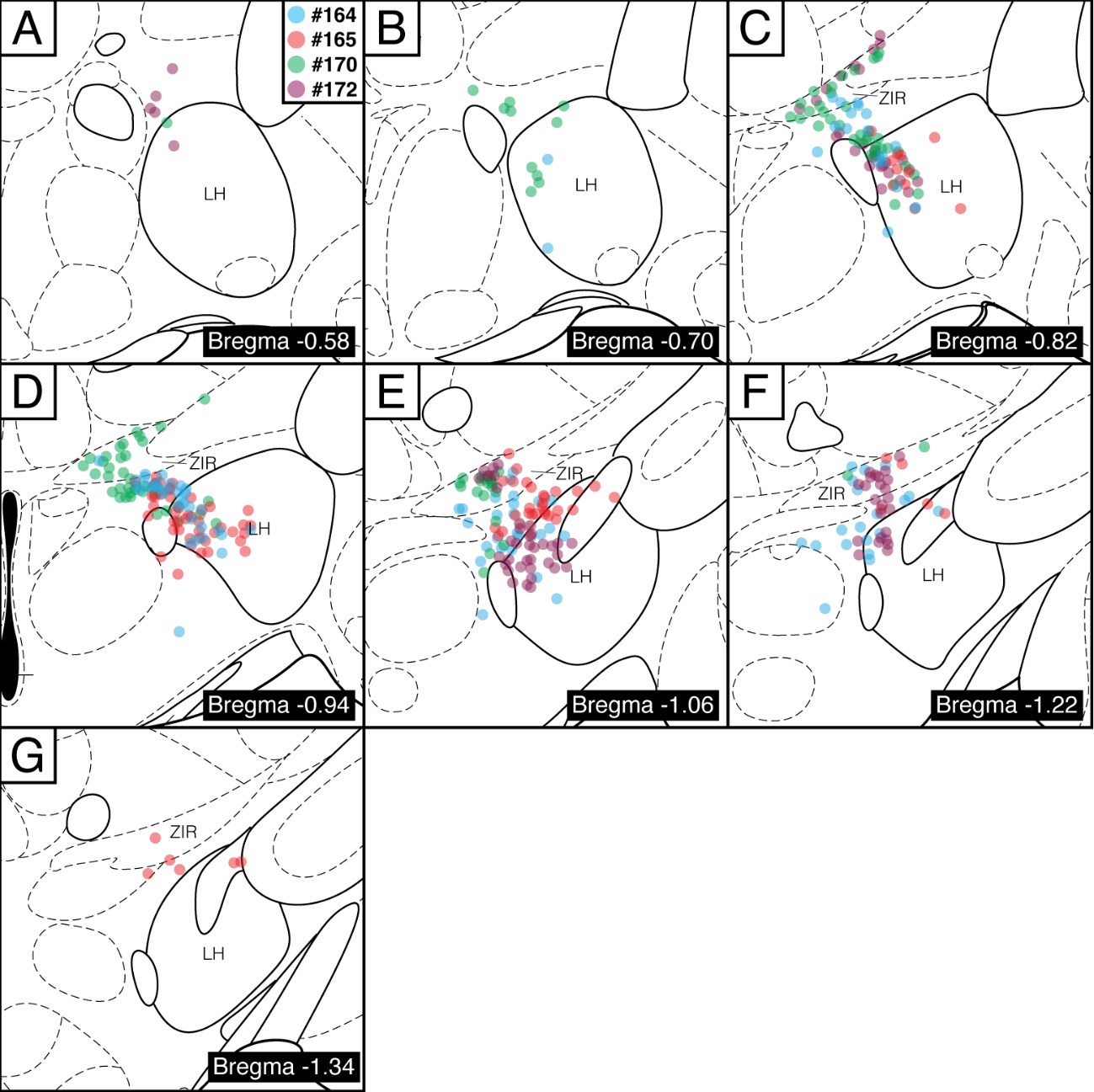

**Supplementary Figure 2.** Coronal brain schematics depicting anterograde tracer injection sites in the ZIR^RXFP3^ group of RXFP3-Cre mice (*n* = 4) mapped onto plates from the Mouse Brain in Stereotaxic Coordinates (Paxinos & Franklin, 2004). Each dot represents an mGFP immunoreactive cell body.

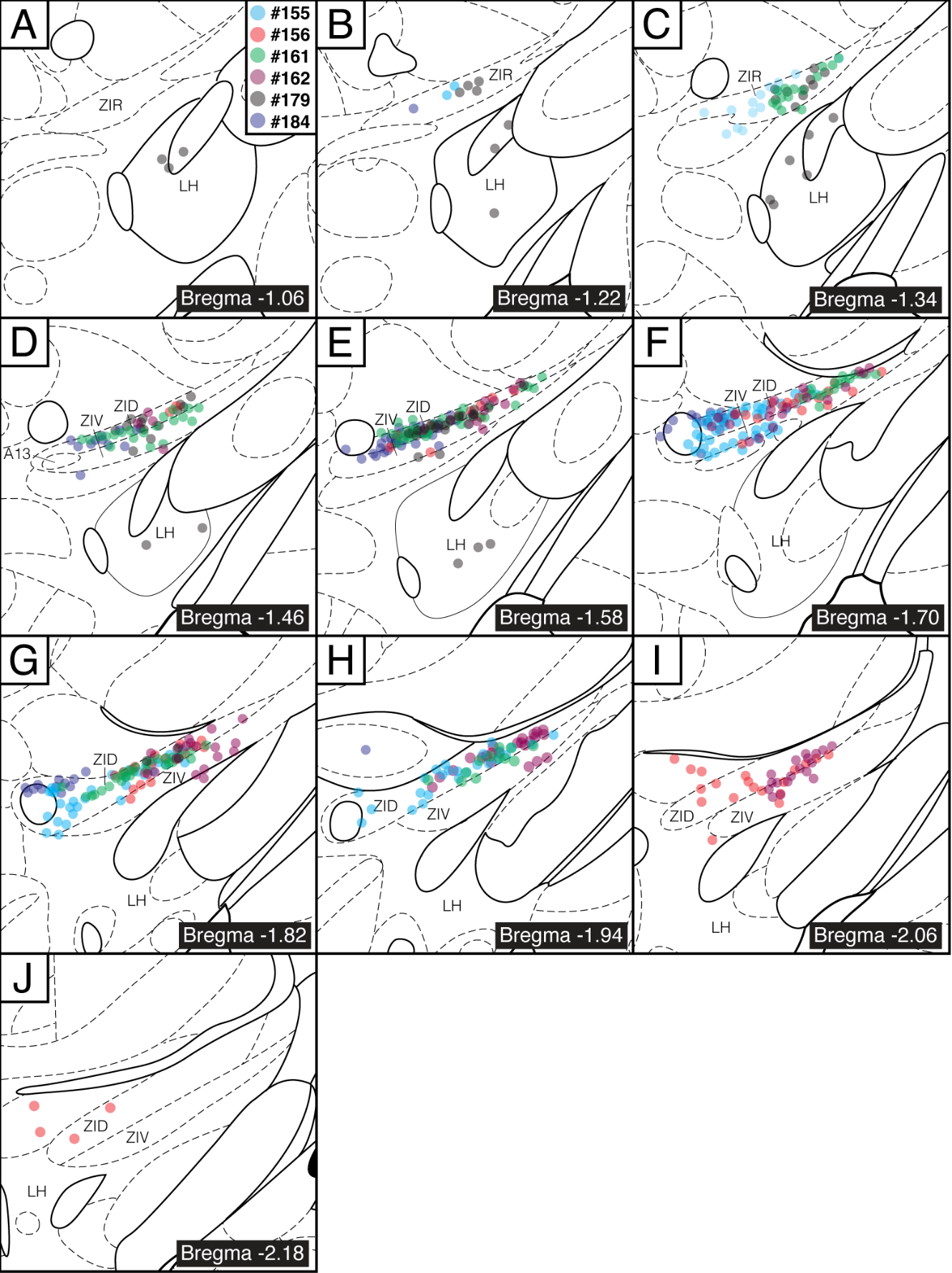

**Supplementary Figure 3.** Coronal brain schematics depicting anterograde tracer injection sites in the ZII^RXFP3^ group of RXFP3-Cre mice (*n* = 6) mapped onto plates from the Mouse Brain in Stereotaxic Coordinates (Paxinos & Franklin, 2004). Each dot represents an mGFP immunoreactive cell body.

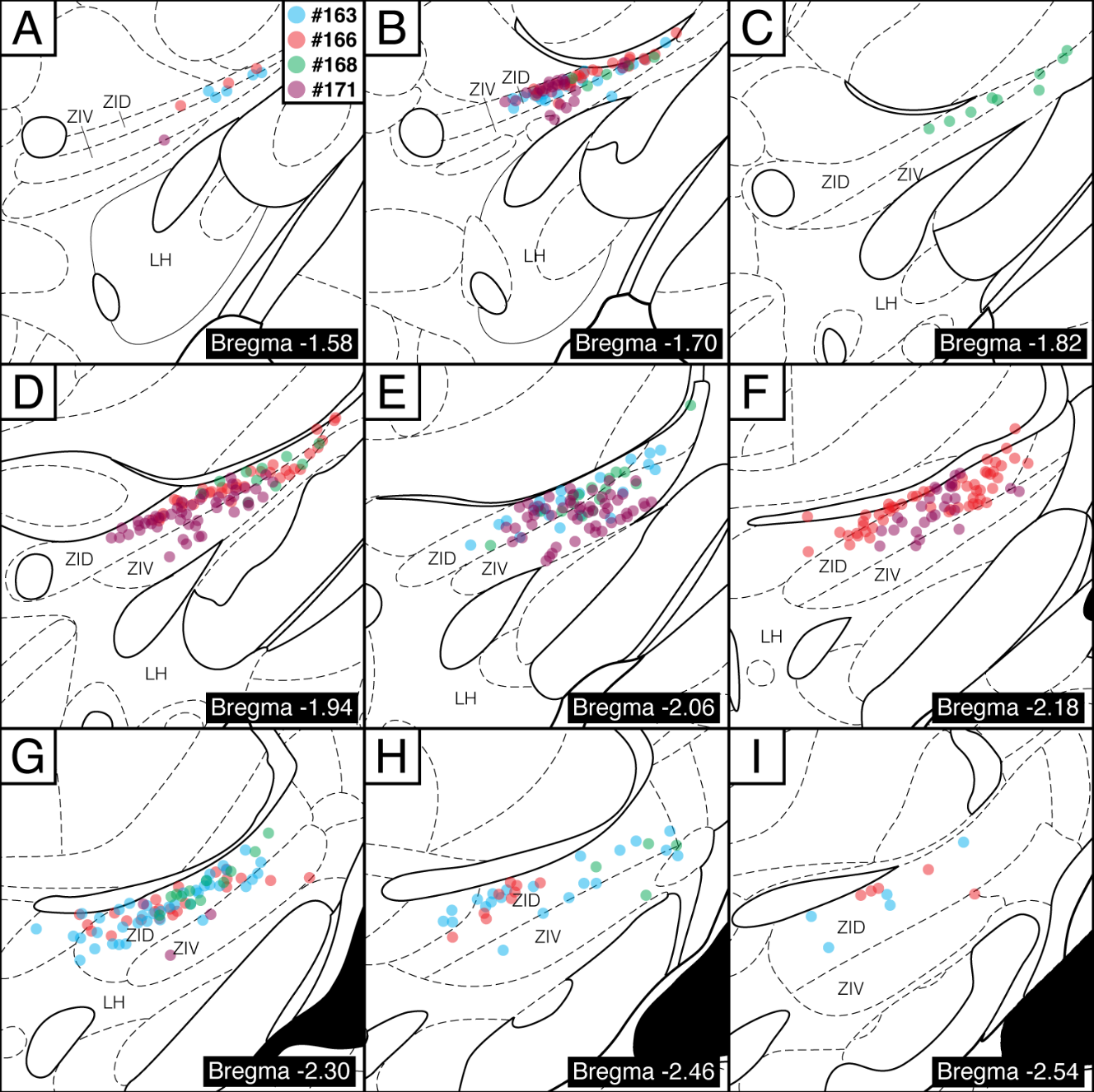

**Supplementary Figure 4.** Coronal brain schematics depicting anterograde tracer injection sites in the ZIC^RXFP3^ group of RXFP3-Cre mice (*n* = 4) mapped onto plates from the Mouse Brain in Stereotaxic Coordinates (Paxinos & Franklin, 2004). Each dot represents an mGFP immunoreactive cell body.

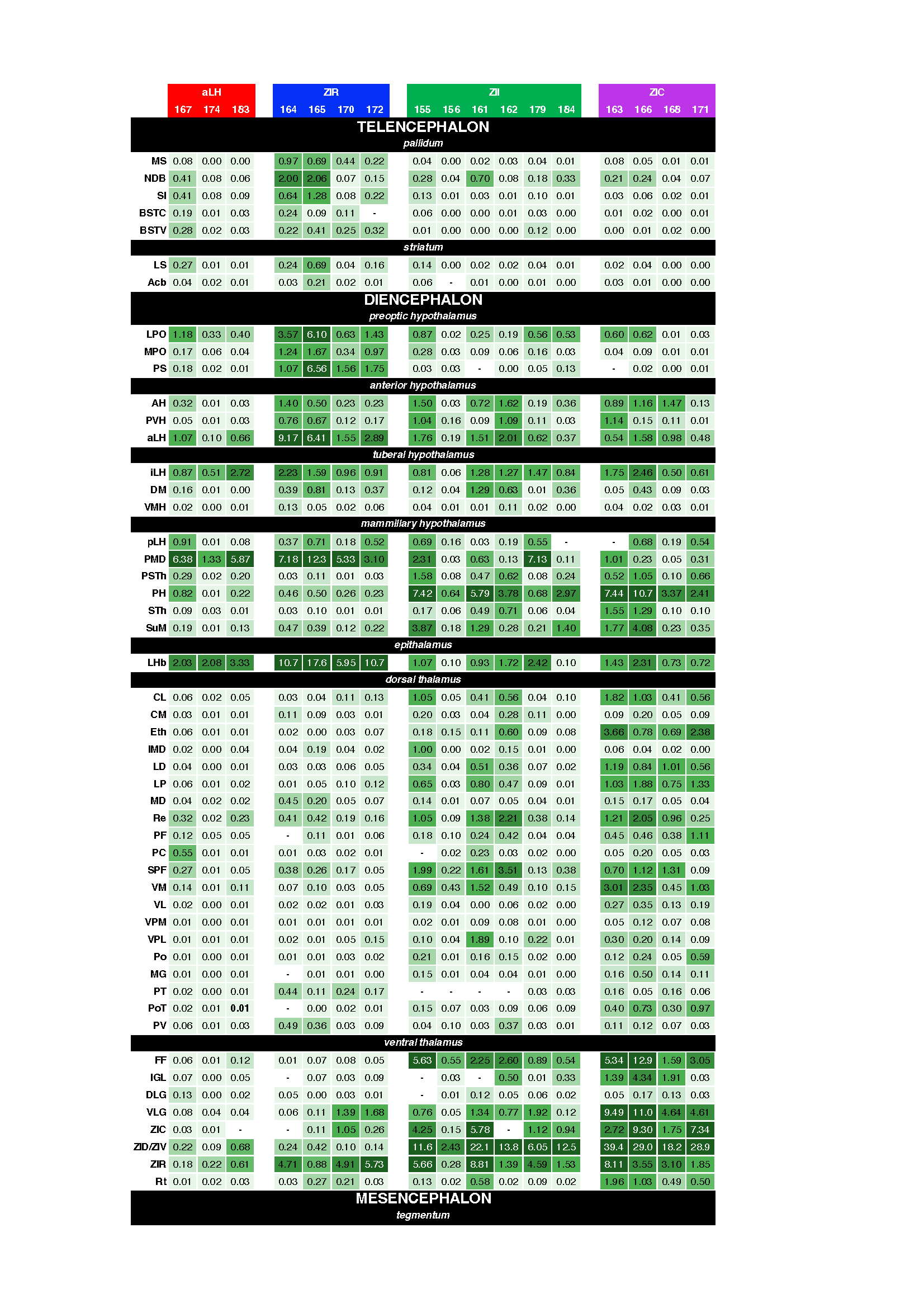

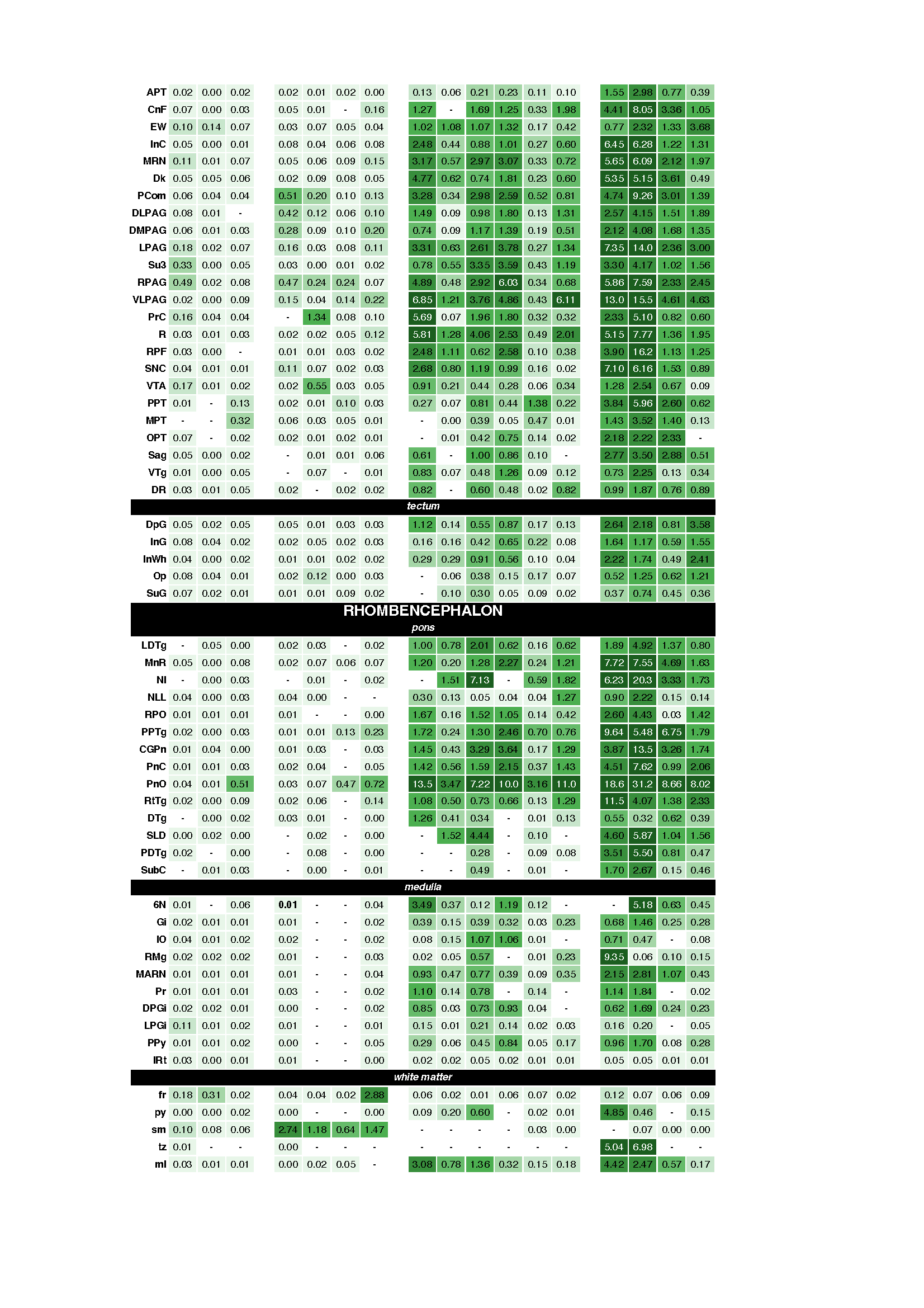
**Supplementary Figure 5.** Heatmap of topographically distinct LH/ZI^RXFP3^ efferent projections for each individual mouse, split by injection site group. Numbers are the average mGFP+ area expressed as a proportion of the total area.

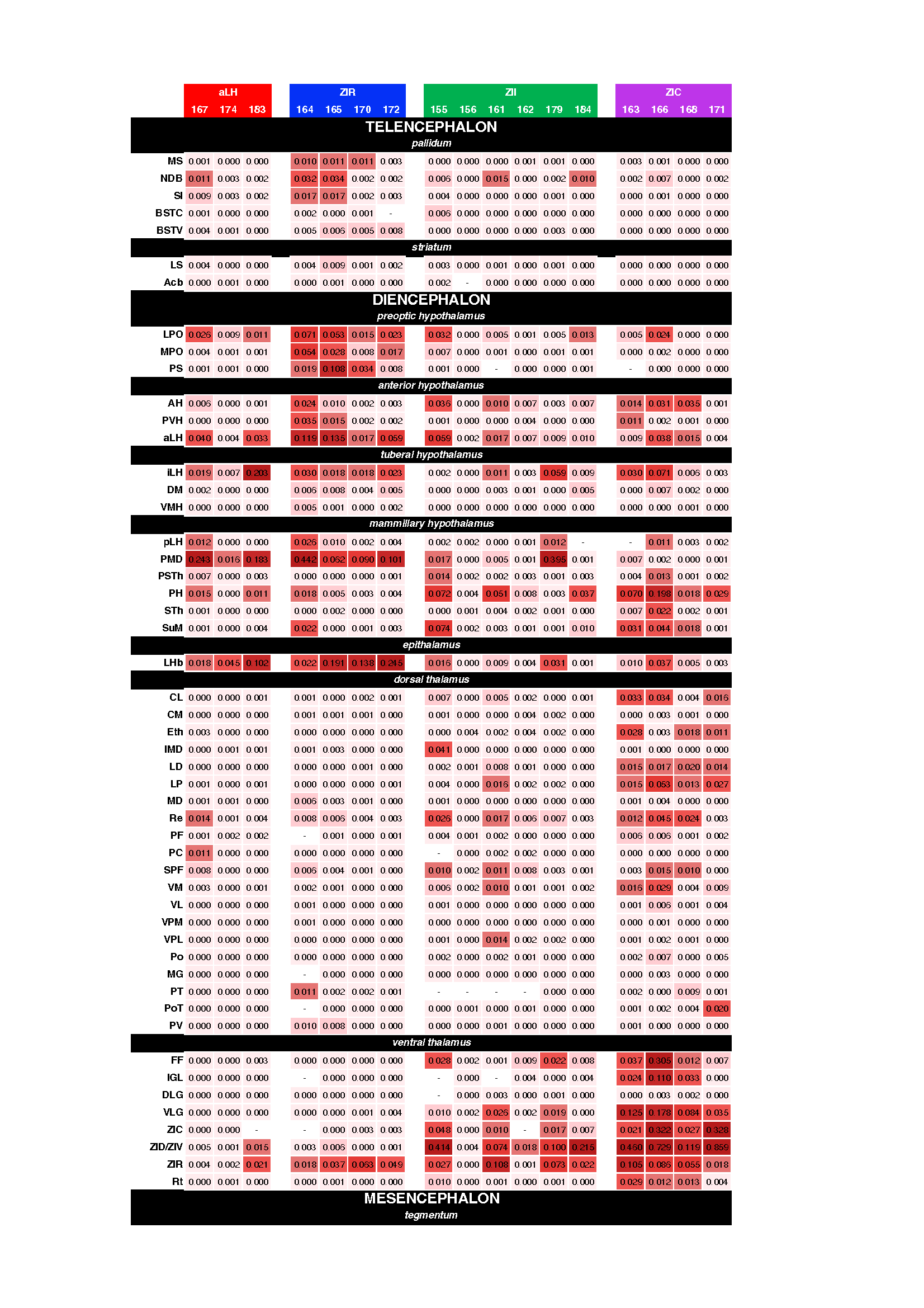

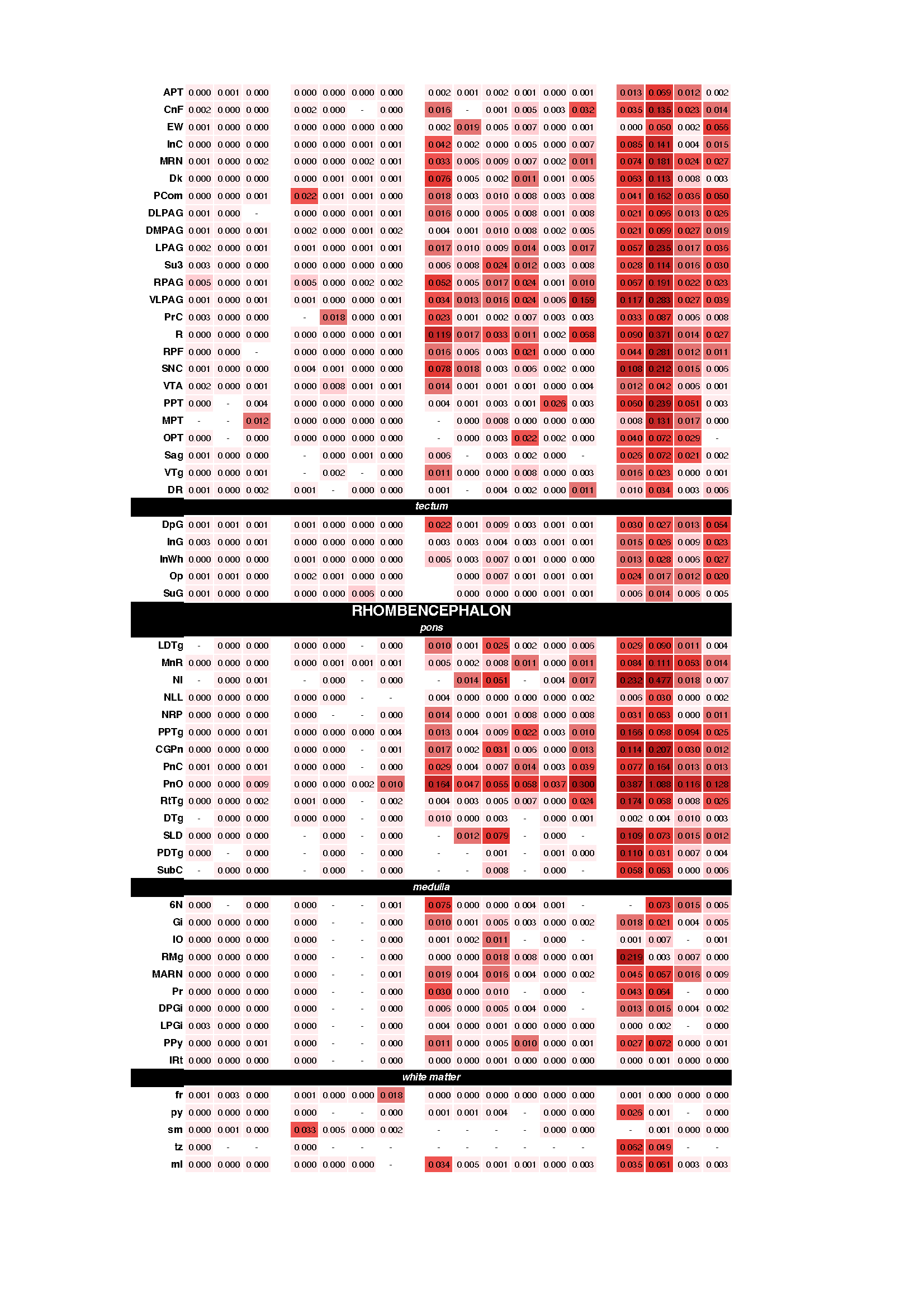

**Supplementary Figure 6.** Heatmap of topographically distinct LH/ZI^RXFP3^ efferent projections for each individual mouse, split by injection site group. Numbers are the average mRuby+ area expressed as a proportion of the total area.

**Supplementary Table 1. Distribution of mGFP+ cells across cases**

| Group | Case | Age | Sex | ZIR | ZID | ZIV | LH | IHy | miss | Total | Range | Notes |
| --- | --- | --- | --- | --- | --- | --- | --- | --- | --- | --- | --- | --- |
| aLH | 167 | 8wk | F | 0 | 0 | 0 | 35 | 1 | 0 | 36 | -0.94 to -1.58 |  |
| aLH | 174 | 8wk | M | 1 | 0 | 1 | 19 | 0 | 0 | 21 | -1.06 to -1.58 |  |
| aLH | 183 | 8wk | F | 3 | 4 | 2 | 37 | 3 | 0 | 49 | -1.06 to -1.70 |  |
| ZIR | 164 | 8wk | F | 16 | 0 | 0 | 36 | 45 | 4 | 101 | -0.70 to -1.22 | 4 mGFP+ cells medial to LH |
| ZIR | 165 | 8wk | F | 11 | 0 | 0 | 52 | 33 | 1 | 97 | -0.82 to -1.34 | 1 mGFP+ cell medial to LH |
| ZIR | 170 | 8wk | F | 43 | 0 | 0 | 26 | 28 | 8 | 105 | -0.58 to -1.22 | 1 mGFP+ cell in Rt, 7 mGFP+ cells outside IHy boundary |
| ZIR | 172 | 8wk | M | 23 | 0 | 0 | 51 | 8 | 8 | 90 | -0.58 to -1.22 | 4 mGFP+ cells in Rt, 4 mGFP+ along Rt/VA border |
| ZII | 179 | 8wk | F | 14 | 29 | 6 | 17 | 1 | 0 | 67 | -1.06 to -1.82 |  |
| ZII | 184 | 8wk | F | 1 | 55 | 11 | 0 | 1 | 2 | 70 | -1.22 to -1.94 | 2 mGFP+ cells in VM |
| ZII | 155 | 12wk | F | 16 | 80 | 21 | 0 | 0 | 0 | 117 | -1.22 to -1.94 |  |
| ZII | 156 | 12wk | F | 0 | 46 | 16 | 0 | 0 | 6 | 68 | -1.46 to -2.18 | 5 mGFP+ cells in VM, 1 mGFP cell medial to ZID boundary |
| ZII | 161 | 10wk | M | 14 | 82 | 23 | 0 | 0 | 3 | 122 | -1.34 to -1.94 | 3 mGFP+ cells in VM |
| ZII | 162 | 10wk | M | 0 | 66 | 30 | 0 | 0 | 0 | 96 | -1.46 to -2.06 |  |
| ZIC | 163 | 8wk | F | 0 | 87 | 18 | 0 | 0 | 8 | 113 | -1.58 to -2.54 | 2 mGFP+ cells in the Rt, 3 mGFP+ cells in the VM, 3 mGFP+ cells in the FF |
| ZIC | 166 | 8wk | F | 0 | 123 | 27 | 0 | 0 | 5 | 155 | -1.58 to -2.54 | 2 mGFP+ cells in the Rt, 3 mGFP+ cells in the VM |
| ZIC | 168 | 8wk | F | 0 | 37 | 18 | 0 | 0 | 1 | 56 | -1.70 to -2.46 | 1 mGFP+ cell in Rt |
| ZIC | 171 | 8wk | M | 0 | 86 | 67 | 0 | 0 | 0 | 153 | -1.58 to -2.30 |  |

|  | **aLH** | | |  | **ZIR** | | | |  | **ZII** | | | | | |  | **ZIC** | | | |
| --- | --- | --- | --- | --- | --- | --- | --- | --- | --- | --- | --- | --- | --- | --- | --- | --- | --- | --- | --- | --- |
| **mGFP+ (%)** | **167** | **174** | **183** |  | **164** | **165** | **170** | **172** |  | **155** | **156** | **161** | **162** | **179** | **184** |  | **163** | **166** | **168** | **171** |
| Pallidum | 8.15 | 4.60 | 1.65 |  | 7.79 | 9.10 | 3.55 | 1.93 |  | 0.17 | 0.07 | 0.26 | 0.05 | 0.92 | 0.17 |  | 0.05 | 0.07 | 0.04 | 0.04 |
| Striatum | 4.78 | 0.86 | 0.24 |  | 1.65 | 5.54 | 0.80 | 1.59 |  | 0.15 | 0.00 | 0.03 | 0.03 | 0.28 | 0.02 |  | 0.02 | 0.02 | 0.01 | 0.02 |
| Hypothalamus | 53.61 | 50.41 | 45.84 |  | 69.31 | 57.42 | 40.76 | 43.09 |  | 10.06 | 4.07 | 9.92 | 9.03 | 18.11 | 5.46 |  | 3.70 | 4.75 | 4.94 | 3.07 |
| Thalamus | 16.53 | 31.89 | 22.16 |  | 15.44 | 22.45 | 33.53 | 30.10 |  | 17.37 | 11.08 | 23.95 | 15.94 | 28.30 | 11.61 |  | 22.06 | 16.03 | 21.91 | 35.38 |
| Midbrain | 12.80 | 5.46 | 6.76 |  | 3.41 | 3.52 | 4.68 | 4.96 |  | 21.94 | 19.60 | 21.04 | 22.19 | 10.27 | 15.19 |  | 20.12 | 23.95 | 18.82 | 20.39 |
| Pons | 1.72 | 2.70 | 12.84 |  | 0.67 | 1.11 | 8.38 | 8.85 |  | 27.04 | 35.85 | 26.62 | 29.81 | 24.17 | 38.93 |  | 33.79 | 34.12 | 34.65 | 24.29 |
| Medulla | 1.44 | 1.33 | 0.47 |  | 0.13 | 0.00 | 0.00 | 0.41 |  | 1.69 | 1.72 | 1.51 | 0.98 | 0.60 | 0.73 |  | 1.52 | 1.74 | 1.11 | 0.78 |
| White Matter | 0.42 | 2.48 | 0.24 |  | 1.37 | 0.39 | 0.82 | 2.73 |  | 0.63 | 0.76 | 0.42 | 0.20 | 0.15 | 0.04 |  | 1.13 | 0.63 | 0.09 | 0.06 |
| **mRuby+ (%)** |  |  |  |  |  |  |  |  |  |  |  |  |  |  |  |  |  |  |  |  |
| Pallidum | 6.95 | 7.34 | 0.87 |  | 5.34 | 10.41 | 6.10 | 2.12 |  | 0.60 | 0.00 | 0.94 | 0.07 | 1.24 | 0.29 |  | 0.07 | 0.06 | 0.01 | 0.04 |
| Striatum | 2.81 | 1.51 | 0.18 |  | 1.31 | 4.62 | 0.61 | 1.47 |  | 0.25 | 0.00 | 0.13 | 0.00 | 0.38 | 0.02 |  | 0.00 | 0.01 | 0.00 | 0.01 |
| Hypothalamus | 69.93 | 46.69 | 67.62 |  | 63.62 | 63.64 | 50.49 | 54.35 |  | 12.75 | 4.47 | 16.61 | 8.05 | 39.83 | 5.16 |  | 3.87 | 4.83 | 6.14 | 1.64 |
| Thalamus | 11.09 | 35.20 | 19.37 |  | 26.49 | 18.51 | 34.85 | 30.61 |  | 31.86 | 4.58 | 24.74 | 9.33 | 34.19 | 11.14 |  | 18.02 | 18.94 | 22.84 | 62.37 |
| Midbrain | 7.22 | 5.20 | 4.54 |  | 2.43 | 2.37 | 4.61 | 2.62 |  | 21.39 | 31.42 | 17.32 | 25.35 | 5.89 | 19.02 |  | 18.57 | 28.45 | 21.50 | 15.26 |
| Pons | 0.85 | 2.21 | 7.23 |  | 0.12 | 0.33 | 3.30 | 7.80 |  | 29.63 | 57.51 | 36.49 | 54.71 | 18.28 | 63.97 |  | 55.93 | 45.72 | 47.52 | 19.93 |
| Medulla | 1.12 | 0.72 | 0.18 |  | 0.01 | 0.00 | 0.00 | 0.28 |  | 2.86 | 1.41 | 3.53 | 2.26 | 0.19 | 0.37 |  | 2.69 | 1.67 | 1.95 | 0.71 |
| White Matter | 0.04 | 1.13 | 0.00 |  | 0.67 | 0.12 | 0.04 | 0.77 |  | 0.66 | 0.60 | 0.24 | 0.22 | 0.00 | 0.03 |  | 0.85 | 0.31 | 0.04 | 0.03 |

**Supplementary Table 2. Percentage of mGFP+ fibres/mRuby boutons observed in each major brain subdivision across cases**
